# Site-specific DNA insertion into the human genome with engineered recombinases

**DOI:** 10.1101/2024.11.01.621560

**Authors:** Alison Fanton, Liam J. Bartie, Juliana Q. Martins, Vincent Q. Tran, Laine Goudy, Matthew G. Durrant, Jingyi Wei, April Pawluk, Silvana Konermann, Alexander Marson, Luke A. Gilbert, Patrick D. Hsu

## Abstract

Technologies for precisely inserting large DNA sequences into the genome are critical for diverse research and therapeutic applications. Large serine recombinases (LSRs) can mediate direct, site-specific genomic integration of multi-kilobase DNA sequences without a pre-installed landing pad, but current approaches suffer from low insertion rates and high off-target activity. Here, we present a comprehensive engineering roadmap for the joint optimization of DNA recombination efficiency and specificity. We combined directed evolution, structural analysis, and computational models to rapidly identify additive mutational combinations. We further enhanced performance through donor DNA optimization and dCas9 fusions, enabling simultaneous target and donor recruitment. Top engineered LSR variants achieved up to 53% integration efficiency and 97% genome-wide specificity at an endogenous human locus, and effectively integrated large DNA cargoes (up to 12 kb tested) for stable expression in challenging cell types, including non-dividing cells, human embryonic stem cells, and primary human T cells. This blueprint for rational engineering of DNA recombinases enables precise genome engineering without the generation of double-stranded breaks.

## Introduction

The prevailing method for a genome insertion edit leverages programmable nucleases for targeted DNA damage, followed by complex interactions with endogenous DNA repair processes, for the final integration of a donor template^1^. The ability to insert multi-kilobase (kb) DNA sequences efficiently and precisely into specified sites in the human genome without generating exposed double-strand breaks or nicks would significantly advance both synthetic biology and gene therapy. This technology could enable the integration of gene circuits and large-scale pooled libraries, as well as the replacement of entire genes rather than the individual correction of diverse patient mutations^2^. However, previous approaches have been hampered by semi-random integration^3^, limited efficiency^4–7^, ceilings on donor template size^8^, or complex multi-component delivery^9–12^.

DNA recombinases are an emerging class of genome editing systems with important mechanistic advantages for achieving precise DNA insertions into the genome. The large serine recombinase (LSR) enzyme family offers high recombination efficiency and site-specificity, operates independently from host repair machinery, and can be delivered as a simple two-component system comprising the recombinase enzyme and the donor DNA^13^. Natively, these enzymes facilitate mobile genetic element integration into bacterial genomes by recombining two double-stranded DNA attachment sites (attP and attB). They are readily adaptable to human cell engineering, integrating at either at pre-installed landing pads or at endogenous pseudosites (termed attH for the human genome) that closely resemble their native integration sequence^13–15^.

Although LSRs have tremendous potential, their broad application is currently constrained by the prerequisite of matching DNA recognition sequences within the target genome. When this sequence is absent, the genome must be pre-engineered to introduce a landing pad – a process particularly challenging in primary cells or *in vivo* settings. Even when suitable pseudosites exist, ensuring high recombinase specificity remains challenging. Our previous screening of over 60 diverse LSR orthologs in human cells revealed that many LSRs integrated into genomic pseudosites, but often with low specificity, targeting hundreds of sites with varying efficiencies. Attempts to modify or engineer LSRs to target endogenous sequences have historically yielded inefficient or non-specific systems^16–18^. Here, we aimed to develop methods to solve the multi-objective optimization problem of high efficiency and high specificity LSRs for direct human genome integration without pre-engineered landing pads.

We reasoned that three of the key mechanistic limitations impeding efficient and specific endogenous integrations are genome recognition and binding, donor DNA binding, and recombination efficiency (**Fig. 1A**). To address these challenges, we developed a comprehensive framework to guide recombinase engineering, using the genome-targeting LSR Dn29 as a proof-of-concept. We employed four distinct strategies: recombinase directed evolution, machine learning-guided mutation stacking, dCas9 fusions, and donor attachment site sequence optimization. Through directed evolution and structural modeling, we generated variants with a 10-fold increase in efficiency and a 5-fold improvement in specificity. Computational models predicted the performance of higher-order combinatorial mutants, streamlining the generation of optimized variants. Modular fusion of LSRs to dCas9 enhanced effector recruitment to the desired genomic pseudosite and donor DNA, and optimized attP sequences further improved donor integration efficiency.

**Figure 1:**
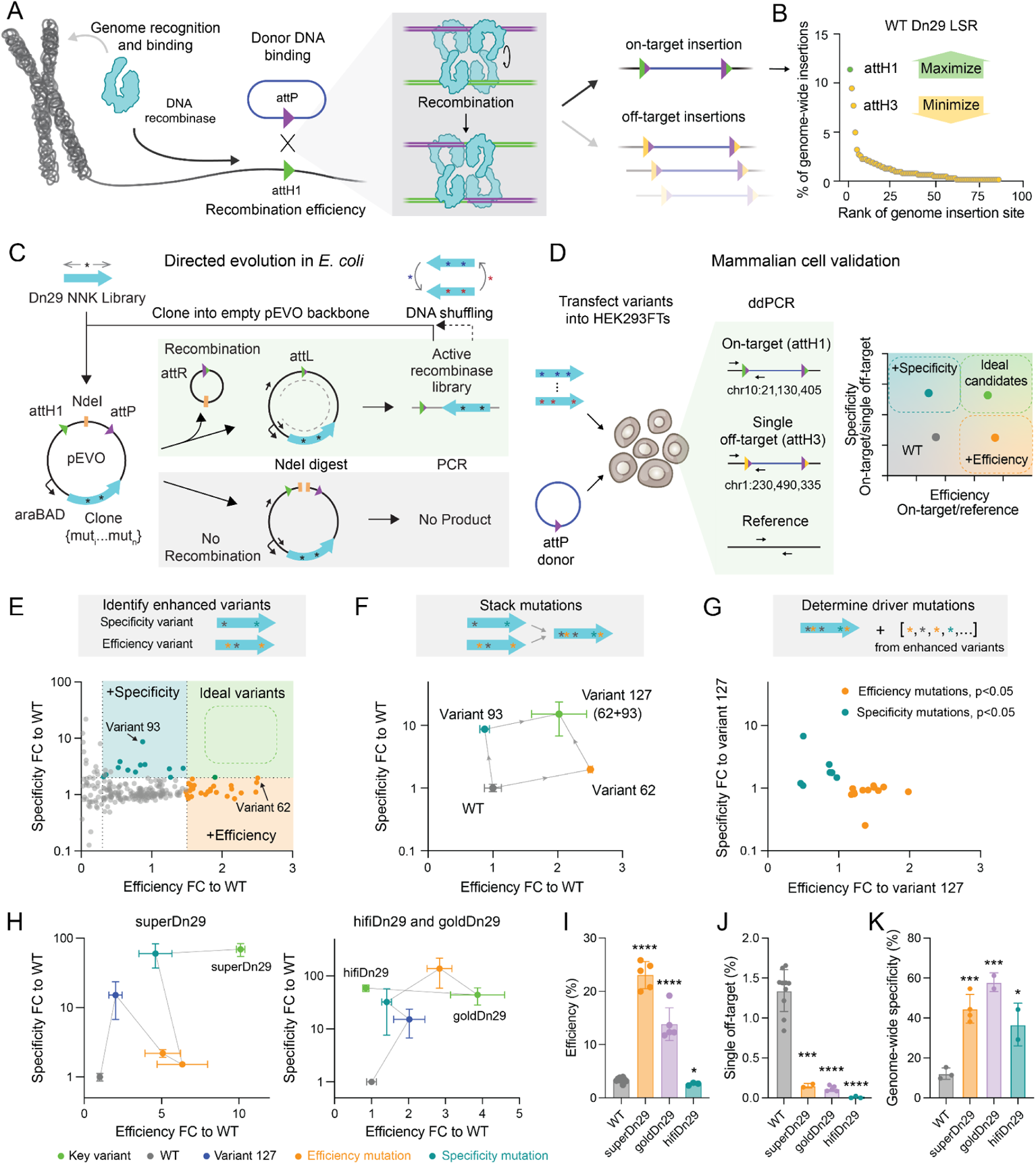
Directed evolution of large serine recombinases (LSRs) with improved efficiency and specificity. A. Overview of engineering strategies to improve integrations into endogenous genomic sites. By improving genome recognition and binding, recombination efficiency, and donor DNA binding, we aim to enhance recombination between a plasmid containing the recombinase attachment site attP and a genomic pseudosite, attH1, by maximizing on-target integrations and minimizing off-targets. B. Genome-wide specificity profile of WT Dn29. The green dot is the on-target site, attH1 (chr10:21,130,405). The yellow dots are off-target sites, ranked by their insertion efficiency. Data shown is the same as Figure 2I. C. Schematic of Dn29 directed evolution scheme in *E. coli*. An evolution backbone, pEVO, expresses a library of Dn29 variants containing NNK codons across the coding sequence and contains attP and attH1 sites. Active variants remove the NdeI site, allowing selective PCR recovery after digestion. The active recombinase library can be shuffled to generate higher order combinations of beneficial mutations and re-cloned into pEVO for subsequent rounds of evolution. D. Schematic of mammalian cell validation of evolved LSR variants. Colonies from the active recombinase library are randomly selected and validated in HEK293FTs. ddPCR at the on-target (attH1), a single-off target (attH3), and a genomic reference measures the efficiency (attH1/reference) and specificity (attH1/attH3). E. Efficiency and specificity of 247 LSR library members in HEK293FTs, shown as fold change (FC) to WT. Colored dots represent enhanced variants with >2-fold WT specificity (teal) or >1.5-fold WT efficiency (orange). Each dot represents the mean of n=2 biological replicates. F. Efficiency and specificity of WT Dn29 (blue, n=16 biological replicates), variant 62 (orange, n=2 biological replicates) and variant 93 (teal, n=2 biological replicates), and variant 127 (green, n=10 biological replicates), generated by stacking all mutations found on variants 62 and 93. Dots and error bars represent the mean ± SD of the biological replicates, shown as FC to WT. G. Efficiency and specificity of significant variants harboring driver mutations (one-tailed p<0.05), shown as FC to Variant 127. Variants are generated by adding individual mutations from the enhanced variants in Figure 1E on top of variant 127. Dots represent the mean of n=2 biological replicates. H. Efficiency and specificity of all variants generated across rounds of mutation stacking, following the path from WT Dn29 (gray, n=16 biological replicates, same data as Figure 1F) to superDn29, goldDn29, and hifiDn29 (green, n=2, 5, and 3 biological replicates, respectively). Variant 127 is shown in blue (n=12 biological replicates, same data as Figure 1F), orange dots indicate the addition of efficiency mutations, and teal dots indicate the addition of specificity mutations, with each dot representing 2-7 biological replicates. Dots and error bars represent the mean ± SD of biological replicates. Gray lines indicate the lineage of mutation addition between variants. I. On-target efficiency of WT (n=10 biological replicates), superDn29 (n=5 biological replicates), goldDn29 (n=5 biological replicates), and hifiDn29 (n=3 biological replicates). Bars and error bars represent the mean ± SD, and dots represent each replicate. Data presented is the same as shown in Figure 1H. Asterisks show t-test significance compared to WT. *=two-tailed p<0.05, ****=two-tailed p<0.0001. J. Efficiency of integration into a single off-target (attH3) of WT (n=10 biological replicates), superDn29 (n=2 biological replicates), goldDn29 (n=5 biological replicates), and hifiDn29 (n=3 biological replicates). Bars and error bars represent the mean ± SD, and dots represent each replicate. Asterisks show t-test significance compared to WT. ***=two-tailed p<0.001, ****=two-tailed p<0.0001. K. Genome-wide specificity of on-target integration compared to all genomic insertions of WT (n=3 biological replicates) superDn29 (n=4 biological replicates), goldDn29 (n=2 biological replicates) and hifiDn29 (n=2 biological replicates). Bars and error bars represent the mean ± SD, and dots represent each replicate. Asterisks show t-test significance compared to WT. *=one-tailed p<0.05, ***=one-tailed p<0.001.

By combining these strategies, we generated a suite of highly efficient recombinases that specifically integrate DNA cargo at a single endogenous locus in the 3.2 billion basepair human genome. Our engineering efforts yielded variants with enhanced characteristics: hifiDn29-dCas9 achieved 97% specificity, superDn29-dCas9 reached 53% efficiency, and our optimized variant, GoldDn29-dCas9, struck a balance between these improvements, attaining up to 90% specificity and 51% efficiency in HEK293FT cells. GoldDn29-dCas9 represents a 7.5-fold improvement in specificity and 12-fold improvement in integration efficiency compared to the wildtype Dn29 enzyme. These enhanced recombinases enabled the site-specific integration of up to 12 kb DNA payloads into various cell types, achieving efficiencies up to 33% in non-dividing cells, 24% in embryonic stem cells, and 17% in primary human T cells. Furthermore, we demonstrated that the targeted integration site maintains strong CRISPRi transgene expression in stem cell-derived hematopoietic progenitor cells, supporting effective knockdown in this differentiated lineage. This work provides a blueprint for rational engineering of recombinases, enabling complex genome engineering processes and permitting easy translation of multi-kilobase genome engineering to new cell types and across model organisms.

## Results

### A framework for recombinase engineering to enable site-specific genome insertion

We previously reported that recombinases can catalyze DNA integration directly into the human genome at a number of endogenous genomic locations, or pseudosites^13^. The number and identity of the pseudosites vary across LSR orthologs based on the sequence similarity between its native attachment site (attB) and the targeted genome (**Fig. 1A**). In this study, we sought to establish the engineering principles that can be used to optimize any LSR for site-specific human genome integration, choosing the genome-targeting LSR Dn29 as a proof-of-concept due to its favorable specificity and efficiency profile^13^. Dn29 integrates into the endogenous genome at a 5% overall efficiency and directs 12% of its insertions into a top site (attH1), with three prominent off-targets (attH2, attH3, and attH4) each comprising 5-10% of total insertions, and ~80 other low frequency off-target sites (**Fig. 1B**).

Our framework involves three key steps: increasing on-target integration efficiency at attH1, reducing insertion frequency at prominent off-targets, and minimizing the long tail of low-frequency off-target insertions. To measure our progress towards these goals, we developed three key metrics: *“efficiency”* as the percentage of attH1 sites that receive an insertion; “*specificity*” as the ratio of insertions into attH1 versus attH3; and “*genome-wide specificity*” as the ratio of attH1 insertions relative to all on- or off-target integration events (**Fig. 1A**, **Extended Data Fig. 1A, Methods**).

### Directed evolution of LSRs with improved efficiency and specificity

Because the protein-DNA recognition code of LSR enzymes for their DNA target site is unknown but unlikely to be modular like zinc finger or TALE proteins, we reasoned that the Dn29 LSR coding sequence would need to be modified via directed evolution to improve on-target integration into attH. To increase Dn29 insertion efficiency at attH1 and disfavor integration into off-targets, we performed deep scanning mutagenesis of Dn29 at single-site saturation and tested the variant library in a intra-plasmid recombination reporter containing the attH1 and attP sites (**Fig. 1C**). Successful recombinants are positively selected by removing an intervening restriction enzyme site, while unproductive variants are eliminated via plasmid digestion^19,20^. To increase overall library complexity beyond single mutants alone, which may not be sufficient for modifying LSR target site preference, we also performed DNA fragmentation and reassembly to shuffle successful mutations (**Extended Data Fig. 1B**)^21^.

After 12 evolution rounds and two DNA fragmentation shuffles, we sequenced the output library using both short and long read-length techniques to leverage the collective benefits of high read depth and the ability to link mutations throughout the full coding sequence of the 519 aa Dn29 LSR (**Extended Data Fig. 1C-1F**). The resulting library contained a median of 3 aa mutations per variant, with 72% of the pool carrying 2-6 mutations (**Extended Data Fig. 1G**). Additionally, the evolution process exhibited robust selection against nonfunctional variants, with mutation dropout rates increasing from 0.0035% in the input library to 44.9% in the output library (**Extended Data Fig. 2A-B**). As expected, we saw strong retention of the catalytic serine residue, depletion of stop codons throughout the Dn29 ORF (**Extended Data Fig. 3A-3B**), and observed that phylogenetic conservation negatively correlated to mutational tolerance (Pearson *r* = −0.3577, two-tailed p<0.0001) (**Extended Data Fig. 3C**). Compared to the input library, the output library was enriched for mutations in several hotspots, particularly in the C-terminal region (**Extended Data Fig. 1E, 3D**). We observed higher mutational sensitivity in the N-terminal domain (NTD), consistent with its important functional roles in inter-subunit interactions, subunit rotation, catalysis, and ligation^22^.

Next, we functionally evaluated variant library members in human cells that were derived from three time points of the directed evolution assay: after 5 cycles, after 7 cycles plus one DNA shuffle, and after 12 cycles plus two DNA shuffles (the final output library). We assessed the efficiency (insertion into attH1) and specificity (attH1/attH3 ratio) metrics of each variant (**Fig. 1D**), observing a significant shift toward higher on-target efficiency between the first and second time points, and more variants with improved specificity after the third time point (**Extended Data Fig. 4A, 4B**).

Because most variants exhibited either enhanced efficiency or specificity, but not both, we hypothesized that combining mutations across these two classes would achieve both desired properties (**Fig. 1E**). First, we combined the four mutations from variant 62 (2.5-fold wild-type (WT) efficiency) and variant 93 (8.6-fold WT specificity) into variant 127, which demonstrated 13.4-fold specificity and 1.8-fold efficiency (**Fig. 1F**).

The ability to simultaneously improve LSR efficiency and specificity motivated us to explore more systematically all mutations in our efficiency- and specificity-enhanced variants. We sought to identify the causal point mutations and remove passenger mutations, thereby enabling higher order mutation stacking. Individual validation of each mutation in variant 62 identified E70G and A224P as the driving efficiency mutations (**Extended Data Fig. 4C**). The sole amino acid mutation in variant 93, N341K, was further investigated by substituting N341 with every amino acid (**Extended Data Fig. 4D**). Many substitutions increased activity, with N341Q improving efficiency 2.3-fold and positively charged residues robustly improving specificity (N341K: 6.7-fold, N341R: 5.3-fold).

We next assessed all point mutations from variants with 1.5-fold WT efficiency (n = 47 mutations) and 2-fold WT specificity (n = 28 mutations). We also included mutations at putative DNA-interfacing residues, chosen based on alignment with the crystal structure of *Listeria* integrase C-terminal domain bound to attP (PDB: 4KIS)^23^, hypothesizing that positively charged mutations could modify DNA binding (**Extended Data Fig. 4E**). These mutations were individually installed into variant 127, identifying 12 additional efficiency and 7 additional specificity driver mutations, each contributing 1.2 to 2.5-fold efficiency and 1.1 to 6.8-fold specificity improvements over variant 127 (**Fig. 1G**). A final round of mutation validation (**Extended Data Fig. 4F**) yielded a final list of 21 efficiency and 12 specificity driver mutations for further rational engineering (**Table S1**).

We optimized Dn29 variants through sequential mutation layering, introducing single mutations into the best variant(s) from the previous round. We began with two lineages: an efficiency lineage containing 341Q and a specificity lineage containing 341K (**Extended Data Fig. 4G**). Double mutation combinations showed mostly additive and subadditive effects, with rare antagonistic or synergistic epistasis (**Extended Data Fig. 4H**). This process yielded three key variants - superDn29 (10-fold efficiency and 70-fold specificity), goldDn29 (4-fold efficiency and 44-fold specificity), and hifiDn29 that combined goldDn29 with four additional specificity driver mutations to achieve WT efficiency with high specificity that approached the ddPCR limit of detection (**Fig. 1H-J, Table S2**). Finally, we conducted whole-genome insertion profiling to confirm the improvement in genome-wide specificity, which increased from 12% with WT Dn29 to 40-60% specificity with the engineered variants (**Fig. 1K**). Overall, our unbiased screening of LSR variants revealed specific mutations driving efficiency or specificity improvements, enabling rational engineering of optimized recombinases.

### Computational modeling and structural analysis of recombinase mutation stacking

We demonstrated that LSR directed evolution followed by classification across our efficiency and specificity metrics enables iterative mutational combination to generate LSRs with a desired functional profile. To expedite experimental testing of larger numbers of variants, we sought to develop a computational model for predicting combinatorial variant activity from single mutant data. The variants were divided into distinct rounds, with the individual mutation validation in round 1 (**Fig. 1G**) and the iterative mutation stacking experiments comprising rounds 2-5. We trained two linear models (linear regression, ridge regression) and two gradient boosting models (XGBoost, CatBoost) on one-hot encoded variant sequences from round 1, and then tested these models on the data from rounds 2-5 (**Fig. 2A**).

**Figure 2:**
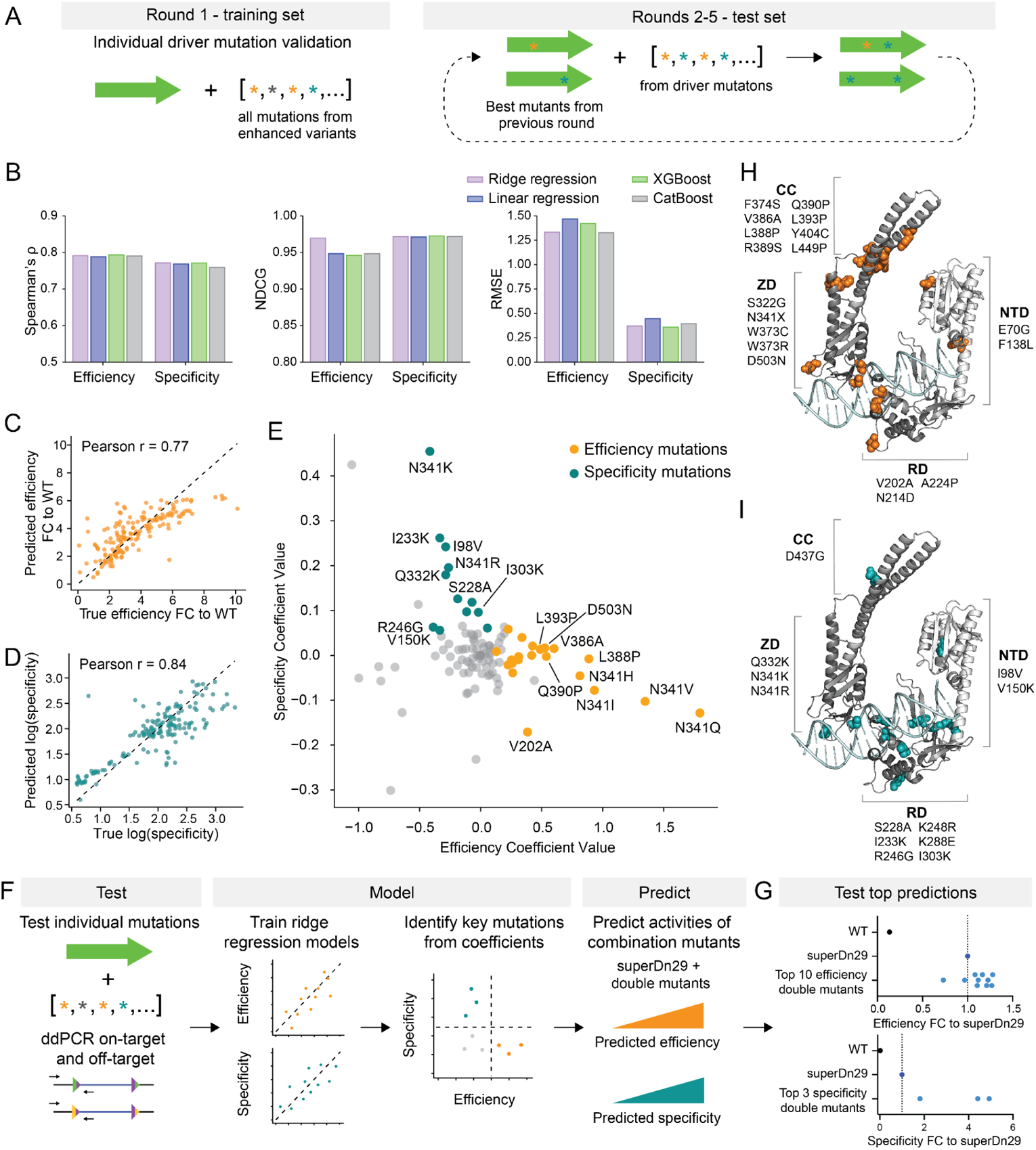
Computational modeling and structural analysis of recombinase mutation stacking. A. Division of data into training and test sets. Models are trained on the individual driver mutation validation data (round 1) and tested on the iterative rounds of higher-order mutants (rounds 2-5). B. Evaluation metrics (Spearman’s *ρ*, normalized discounted cumulative gain (NDCG), root-mean squared error(RMSE)) of linear regression, ridge regression, XGBoost, and CatBoost models. C. Predicted versus true efficiency of higher-order mutants, as fold change (FC) to WT. Pearson r = 0.77, two-tailed p<0.0001. D. Predicted versus true specificity of higher-order mutants, as the log transformation of the ratio of attH1/attH3. Pearson r = 0.84, two tailed p<0.0001. E. Coefficient values of mutations in the efficiency and specificity models. Colored dots indicate mutations identified as efficiency (orange) or specificity (teal) driver mutations from Figure 1. F. Schematic of the workflow for testing and modeling single mutations for the prediction of protein activities of higher-order mutants. G. Efficiency (top) and specificity (bottom) of top model-guided combinatorial mutants, generated by predicting the activity of combining two mutations on top of superDn29. Each dot represents the mean of n=6 biological replicates for WT and superDn29, and n=2 biological replicates for all model-guided variants. H-I. (H) Efficiency and (I) specificity mutations mapped to the Alphafold3 structure of Dn29 bound to attB-R. NTD: N-terminal domain; RD: recombinase domain; ZD: zinc-ribbon domain; CC: coiled-coil motif.

We evaluated model performance using root-mean squared error (RMSE), Spearman’s rank correlation, and normalized discounted cumulative gain (NDCG). NDCG was prioritized as the key evaluation metric because it emphasizes higher-activity mutants, matching our experimental validation priorities. While all models performed well, ridge regression stood out for both efficiency and specificity, showing high accuracy (NDCG = 0.970 for efficiency, 0.971 for specificity) in predicting higher-order mutant activities (**Fig. 2B-D**). The performance of linear models in extrapolating single mutant activity to higher-order mutants supports our previous observations (**Extended Data Fig. 4H**) that LSR mutations are largely additive, and aligns with previous research showing that regularized linear models perform well across mutagenesis datasets of 14 different enzymes^24^.

To identify key mutations, we examined regression coefficients, revealing strong concordance between computationally and experimentally identified impactful mutations (**Fig. 2E**, **Extended Data Fig. 5A-B**). The model coefficients strongly correlated with experimental fold changes (efficiency: ρ=0.623, two-tailed p=0.0025; specificity: ρ=0.930, two-tailed p<0.0001; **Extended Data Fig. 5C-D**), demonstrating that the interpretability of linear models enables the automated identification of impactful mutations.

We outline an approach for applying regression models to predict the efficiency and specificity of higher order mutants (**Fig. 2F**). From a dataset of individual mutations, measured for specificity and efficiency by ddPCR, we generated ridge regression models and examined their coefficients to quantify the impact of key mutations. By predicting efficiency and specificity activity of combinations of key mutations, we can prioritize mutants for experimental testing. To demonstrate this approach, we predicted the activities of all double mutants combined on top of superDn29, skipping round 6 to directly test round 7 of iteration. We tested the top 10 efficiency and top 3 specificity variants for genome integration, and found that 8/10 efficiency variants and 3/3 specificity variants performed better than superDn29 (**Fig. 2G**, **Extended Data Fig. 5E-G**). Overall, we demonstrate that model-guided recombinase design across multiple activity axes can be used to further push efficiency and specificity beyond a space that is easily reachable by traditional directed evolution.

To gain mechanistic insights from our mutational landscape, we generated an AlphaFold3 structural model of Dn29 bound to the attB-R DNA half-site (**Fig. 2H-I**, **Extended Data Fig. 6A**)^25^, which showed high alignment (Zinc-ribbon domain RMSD = 1.341, Recombinase domain RMSD = 1.707) to the *Listeria* integrase C-terminal domain bound to attP (PDB: 4KIS)^23^ (**Extended Data Fig. 6B**). We identified an efficiency mutation hotspot (residues 373-393 and 449) in the coiled-coil domain hinge region, potentially modifying tetramer stabilization or autoinhibitory control (**Extended Data Fig. 6C**)^26^. Despite previous work on a different ortholog that identified mutations in this region as enabling the excision reaction^27^, our key variants maintained unidirectionality (**Extended Data Fig. 6D**). Another key efficiency mutation (D503N) reduces negative charge in a tri-aspartic acid stretch (503-505), likely enhancing DNA phosphate backbone interactions (**Extended Data Fig. 6E**).

Many specificity driver mutations localize near the DNA-binding interface, often replacing neutral amino acids with positively charged ones, potentially strengthening DNA interactions (**Extended Data Fig. 6F**). We also observed efficiency driver mutations near the DNA-interfacing regions, which may modify protein conformation and influence DNA affinity. This combined computational and structural analysis provides a multifaceted understanding of how specific mutations impact LSR function, informing rational engineering of optimized variants to enhance LSR performance.

### Programmable target and donor DNA recruitment with LSR-dCas9 fusions

We reasoned that the LSR protein and target DNA interaction could be further enhanced by developing programmable LSR-dCas9 fusions that facilitate LSR recruitment to the genomic target site. We first optimized the fusion design, observing that an N-terminal dCas9 fusion abolished Dn29 activity, likely due to steric hindrance of tetramerization as the N-terminus is located at the tetrameric complex core^23^. In contrast, C-terminal fusions supported robust recombination and were used for further experiments (**Fig. 3A**).

**Figure 3:**
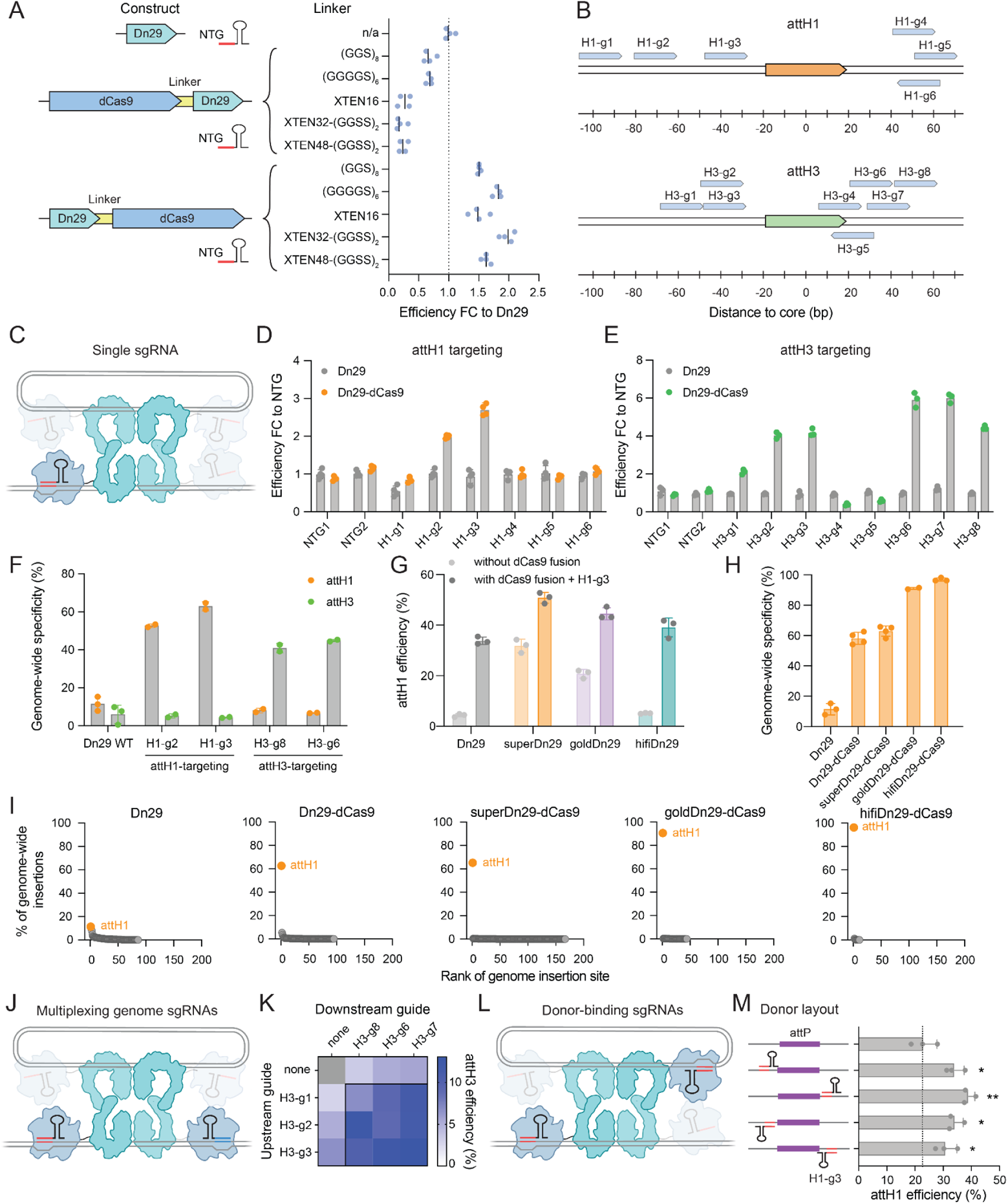
Programmable target and donor DNA recruitment with LSR-dCas9 fusions. A. Schematic of LSR-dCas9 fusion orientations and recombination efficiency with various linkers at attH1 with a non-targeting guide (NTG). The lines represent the mean of n=4 biological replicates, shown as dots. B. Schematic of sgRNA targets for the attH1 (chr10:21,130,405) and attH3 (chr1:230,490,334) pseudosites. C. Schematic of the LSR-dCas9 tetrameric complex, with a single sgRNA targeting the genome. D-E. Integration efficiencies of Dn29 and Dn29-dCas9 at (D) attH1 and (E) attH3 with sgRNAs targeting proximal to the respective pseudosites. Data shown as fold change relative to non-targeting guide (NTG). Bars and error bars represent mean ± SD of n = 3 biological replicates, shown as dots. F. Biasing Dn29-dCas9 integrations to different pseudosites. Shown is the percent of all genome-wide insertions at attH1 (orange) and attH3 (green), using sgRNAs targeting the pseudosites. The bars and error bars represent the mean ± SD of n=3 or 2 biological replicates, shown as dots. G. Integration efficiencies of Dn29 variants at attH1, with and without the dCas9 fusion and guide H1-g3. The bars and error bars represent the mean ± SD of n=3 biological replicates, shown as dots. Data shown is the same as presented in Figure 5B. H. Genome-wide specificity to attH1 of Dn29 and key variants fused to dCas9, targeting attH1 with H1-g3. Bars and error bars represent the mean ± SD of n=2-4 biological replicates, shown as dots. Data shown combines replicates transfected with WT attP and e-attP. I. Representative replicates of genome-wide specificity profiles of Dn29 and key variants fused to dCas9. Orange dots represent the on-target locus (attH1) and gray dots represent off-target loci. Data shown for Dn29 is the same as presented in Figure 1B. J. Schematic of the LSR-dCas9 tetrameric complex, with multiplexed sgRNAs targeting the genome upstream and downstream of the pseudosite. K. Heatmap showing attH3 integration efficiencies (%) of Dn29-dCas9 using guides targeting upstream and downstream of the pseudosite, individually and multiplexed. Each cell represents the mean of n=3 biological replicates. L. Schematic of the LSR-dCas9 tetrameric complex with a single sgRNA targeting both the genome and the donor plasmid. M. Integration efficiencies of Dn29-dCas9 using donor plasmids with the H1-g3 sgRNA target sequence adjacent to the attP. Plasmid schematics show sgRNA target placement (5’ or 3’, top or bottom strand). The bars and error bars represent the mean + SD of n=3 biological replicates, shown as dots. Asterisks show t-test significance compared to wildtype donor plasmid. *=one-tailed p<0.05, **=one-tailed p<0.01.

We next evaluated the impact of dCas9-mediated genomic recruitment using sgRNAs targeting regions near attH1 or attH3 (**Fig. 3B, C**). This single-guide approach increased integration efficiency 2.7-fold at attH1 and 6-fold at attH3 compared to non-targeting guides (**Fig. 3D, E**). We further improved efficiency by optimizing the ratio of donor, effector, and guide components (**Extended Data Fig. 7A**), and confirmed the importance of direct tethering of Dn29 to the genomic target site, as replacement of the linker with a 2A peptide to induce ribosomal skipping abolished the effect (**Extended Data Fig. 7B**).

Interestingly, we could manipulate Dn29’s natural integration preference using dCas9 recruitment. While WT Dn29 naturally integrates into attH1 with 2-fold frequency over attH3, dCas9-based recruitment to attH1 amplified this preference to 14-fold. Conversely, attH3-targeting sgRNAs reversed this bias, resulting in 11-fold higher integration at attH3 compared to attH1 (**Fig. 3F**). Combining the dCas9 fusion strategy with our optimized Dn29 variants further improved on-target efficiency. SuperDn29-dCas9 achieved 50.8% integration at attH1, while goldDn29-dCas9 and hifiDn29-dCas9 reached 44.5% and 39.1% respectively, representing up to an 11.8-fold efficiency increase (**Fig. 3G**).

Next, we tested if dCas9 fusion enhanced LSR specificity by biasing insertion towards the desired pseudosite. Whole-genome insertion profiling showed on-target integrations improved from 12% with WT Dn29 to over 60% with Dn29-dCas9 targeted to attH1, though rare off-target sites persisted. The superDn29-dCas9 variant maintained a similar insertion profile but moderately increased rare off-target sites due to higher overall activity. Notably, goldDn29-dCas9 and hifiDn29-dCas9 achieved 91% and 97% genome-wide specificity to attH1, with significantly fewer off-target loci (35 and 12 sites, on average, respectively) (**Fig. 3H, I**). We also examined the consistency of off-target events across replicates. For Dn29-dCas9, 20-35% of off-target sites were shared between replicates. This decreased to 6-12% for superDn29-dCas9, and to 0% for goldDn29-dCas9 and hifiDn29-dCas9 (**Table S3**), demonstrating robust off-target elimination.

We assessed the generalizability of our fusion approach across 3 additional LSR orthologs (the genome-targeting LSRs Pf80 and Nm60, and the landing pad LSR Si74), demonstrating up to 10-fold improved efficiency (**Extended Data Fig. 7C-7F**, **8A-8C**). To elucidate the parameters of optimal sgRNA design, we analyzed integration efficiencies across all fusion variants and orthologs and determined optimal sgRNA placement to be ~40 bp from the attachment site core, agnostic of guide orientation (**Extended Data Fig. 8D**).

We next explored three orthogonal strategies to further improve efficiency, using dCas9 variants with alternative PAM recognition^28^, multiplexing sgRNAs, and incorporating sgRNA binding sites on donor plasmids. dCas9 variants with increased PAM flexibility can improve efficiency by expanding pseudosite-proximal guide options and enabling more precise sgRNA placement. Using dCas9-HF1-SpG (NGN PAM), we achieved 22% insertion efficiency at the best NGH guide compared to 15% with the best NGG guide (**Extended Data Fig. 8E-8G**). Multiplexed sgRNAs targeting upstream and downstream of the attachment site could further improve genome search and binding (**Fig. 3J**). At the attH3 site, the H3-g3 and H3-g7 combination achieved 13% integration versus 6.6% and 5.6% efficiencies individually (**Fig. 3K**). Finally, we tested the ability of an sgRNA-binding sequence on the donor plasmid to facilitate donor recruitment, across four orientations flanking the minimal attP site (top and bottom strand, upstream and downstream) (**Fig. 3L**). All four configurations improved integration efficiency for Dn29-dCas9, reaching up to 40% integration compared to 23% with the WT donor (**Fig. 3M**). Similarly, incorporating the Nm60-g1 sequence on the donor plasmid improved Nm60 attH2 integration from 61% to 73% (**Extended Data Fig. 8H**).

Our results show that LSR-dCas9 fusions can substantially enhance both efficiency and specificity of cargo DNA insertion by improving LSR recruitment to target and donor DNA. The positive correlation between efficiency and specificity (**Extended Data Fig. 8I**) suggests that optimizing guide RNA sequences and employing effective multiplexing strategies are crucial for maximizing both parameters simultaneously. This approach, combined with engineered LSR variants and dual-targeting guides, achieves up to 97% specificity or over 73% efficiency at a single genomic locus.

### Exploring attP sequence space enables the design of optimized donor DNA

Beyond the use of dCas9 fusions, we explored enhancing donor DNA recruitment by optimizing the attP sequence. We created two Dn29 attP donor plasmid libraries, each with one constant half-site and the other mutated using custom mixed base oligos for an average of ~5.5 mutations per 26 bp half site (**Methods**). Transfecting these libraries into cells expressing WT Dn29-dCas9 and H1-g3 guide, we sequenced attH1 integrants and calculated nucleotide enrichment scores (**Fig. 4A-C**). While the WT nucleotide was generally preferred, we identified 6 positions where non-WT nucleotides showed moderately higher enrichment. Combining all 6 substitutions into an optimized e-attP sequence improved integration efficiency by 1.3-fold. Applying this strategy to Nm60 with Nm60-dCas9 and Nm60-g2, we identified an e-attP with 7 mutations that improved efficiency by 1.1-fold (**Fig. 4E, 4F**).

**Figure 4:**
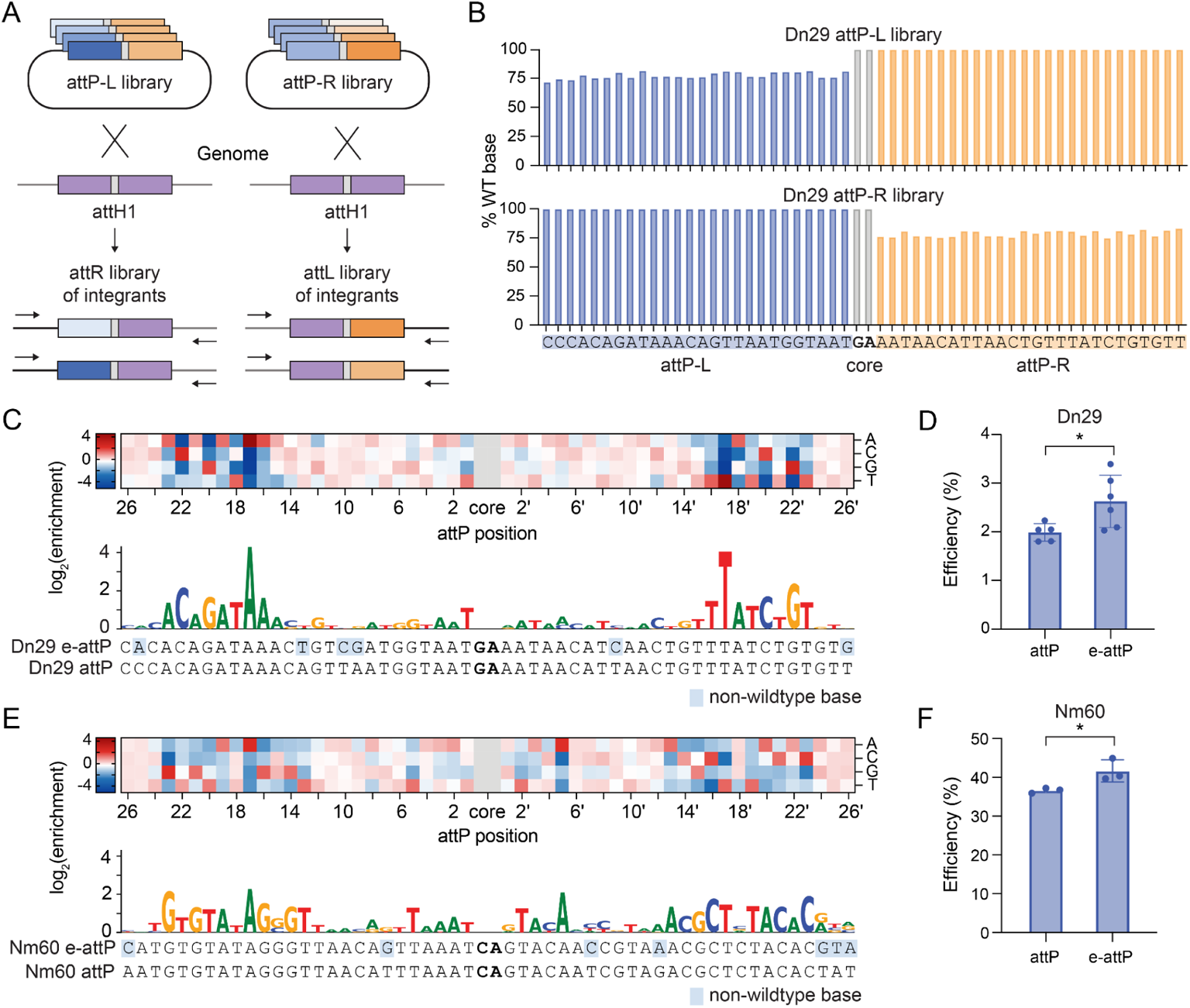
Exploring attP sequence space enables the design of optimized donor DNA. A. Schematic of attP optimization screen. attP-L and attP-R libraries are transfected into HEK293FTs with Dn29-dCas9, and integrants are sequenced. B. Design of attP-L and attP-R libraries. Libraries are generated using mixed base oligos containing 79% WT and 7% each other nucleotide for one half site and constant WT sequence for the other half site. C. Dn29 attP nucleotide enrichment/depletion heat map (top) and sequence logo of enriched nucleotides in e-attP (bottom). Data represent average enrichment scores of two biological replicates. D. Dn29 integration efficiency with attP and e-attP donor plasmids. The bars and error bars represent the mean ± SD of n=5 (attP) and n=6 (e-attP) biological replicates, shown as dots. Asterisks show t-test significance compared to wildtype donor. *= one-tailed p<0.05. E. Nm60 attP nucleotide enrichment/depletion heat map (top) and sequence logo of enriched nucleotides in e-attP (bottom). Data represent average enrichment scores of two biological replicates. F. Nm60 integration efficiency with attP and e-attP donor plasmids. The bars and error bars represent the mean ± SD of n=3 biological replicates, shown as dots. Asterisks show t-test significance compared to wildtype donor. *= one-tailed p<0.05.

Our library enrichment approach provides a high-resolution view of DNA specificity and recombination efficiency determinants. For both Dn29 and Nm60, native attachment sites appear highly evolutionarily optimized, with only a few positions showing incremental efficiency improvements through mutation.

We identified core-distal regions of outsized importance for functional recombination: positions 16-23 and 16’-23’ for Dn29 (**Fig. 4C**) and 13-23 and 13’-23’ for Nm60 (**Fig. 4D**), consistent with previous reports on other LSR orthologs^29–31^. Dn29’s position 17/17’ showed particularly high enrichment, strongly preferring A and T respectively. Notably, the number of highly enriched core-proximal positions varies significantly between orthologs: 1 for Dn29, 3 for Nm60, and 10 for the LSR ortholog Bxb1^29^. These attP studies indicate that donor DNA recognition sequences should be optimized for each individual ortholog, although larger-scale studies may eventually reveal more predictive patterns.

While LSR attachment sites are canonically imperfect inverted repeats, the significance of this asymmetry remains unclear^31^. Our data reveals that the most strongly enriched nucleotides are symmetrical across the core, aligning with previous Bxb1 studies^29,32^. However, for Dn29 and Nm60, we observed 5 and 8 nucleotides with subtle preferences for asymmetric nucleotides at corresponding half-site positions. This suggests that slight attachment site asymmetry may be a deliberate feature of the recombination mechanism, rather than a consequence of mutations or phage genome sequence constraints. Overall, this attachment site exploration deepens our understanding of LSR target site recognition and advances our ability to design optimal DNA donors.

### Unifying engineering strategies for maximal LSR efficiency and specificity

Finally, we aimed to create optimal LSR tools for large DNA cargo integration by combining our orthogonal engineering efforts. Armed with directed evolution variants, dCas9 fusions with sgRNA design rules, and optimized donor DNA substrates, we assessed the effects of combining these features into a single system (**Fig. 5A-B**).

**Figure 5:**
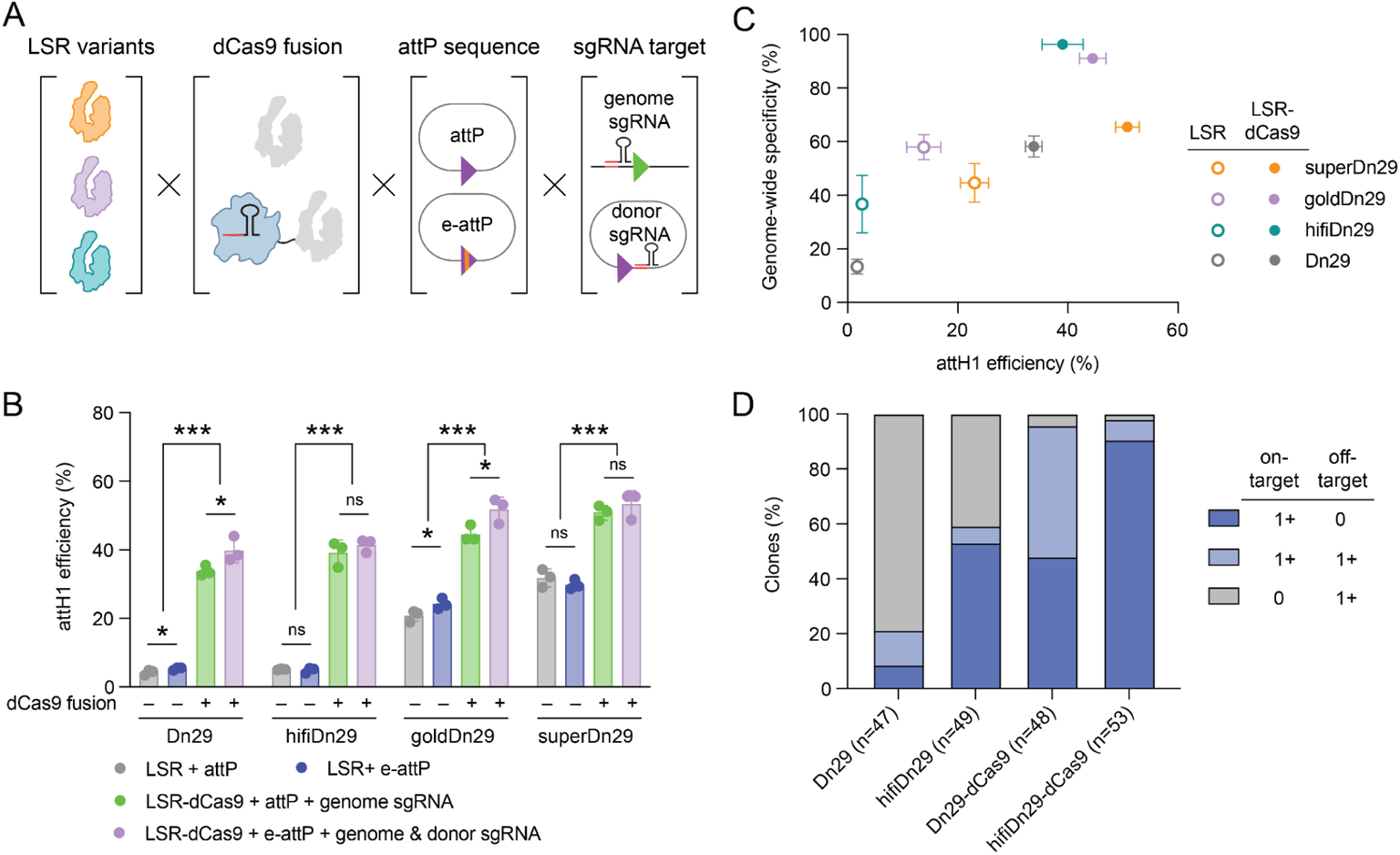
Unifying engineering strategies for maximal LSR efficiency and specificity. A. Schematic overview of developed engineering strategies. B. Integration efficiencies at attH1 of Dn29 with combined engineering strategies. The bars and error bars represent the mean ± SD of n=3 biological replicates, shown as dots. Asterisks show t-test significance. * = one-tailed p<0.05;***=one-tailed p<0.001; ns=not significant. C. Integration efficiency and genome-wide specificity of engineered LSRs at attH1, with and without dCas9 fusions and e-attP/donor sgRNA donors. Dots and error bars represent the mean ± SD of n=2 biological replicates. D. Specificity analysis of HEK293FT single cell clones engineered with Dn29 and hifiDn29, with and without dCas9 fusions. N of clones analyzed per sample is indicated in the x-axis.

Overall, our combined engineering strategies substantially improved recombination efficiency; the variants fused to dCas9 with the optimized donor achieved 41-53% efficiency, a 9.6- to 12.3-fold improvement over the WT enzyme (**Fig. 5B, Table S4**). The relative impact of combining attP optimization and dCas9 fusion with Dn29 or its variants depended on the baseline efficiency of each recombinase. Given the relatively higher efficiency of goldDn29 and superDn29, the improvements were more modest (1.6 to 2.1-fold) compared to Dn29 and hifiDn29 (7.6 to 7.8-fold). Interestingly, the e-attP did not significantly improve integration efficiency for superDn29 or hifiDn29, indicating future variant-specific attP optimization may be beneficial.

Using hifiDn29-dCas9, our most specific configuration, we measured 97% genome-wide specificity by bulk integration site sequencing (**Fig. 5C**). To better understand the single cell variation of insertional mutagenesis, including on-/off-target co-occurrence and integration copy number, we mapped integrations in ~50 clonal HEK293FT populations edited with hifiDn29 or WT Dn29, with and without dCas9 fusions (**Fig. 5D**). dCas9 fusion dramatically improved performance for both Dn29 and hifiDn29, resulting in over 95% of clones containing on-target edits. However, the most striking difference was observed in off-target insertions - 91% of Dn29 clones contained off-target insertions, compared to 46% of hifiDn29 clones. Ultimately, with dCas9 fused to hifiDn29, off-target insertions were reduced to only 9% of clones, compared to 52% for dCas9-Dn29. These single cell specificity measurements closely mirrored the bulk genome-wide specificity results, validating the accuracy and sensitivity of the genome-wide specificity assay (**Extended Data Fig. 9A**).

Beyond targeting accuracy, measuring the number of integrations per cell is crucial for fully understanding editing outcomes. Because dCas9 increases efficiency, it also increases the rate of multiple on-target insertions. HifiDn29 showed the highest rate of single on-target insertion events, with half of the clones exhibiting this genotype. In contrast, including the dCas9 fusion decreased the rate of single on-target insertion events to 38%, and increased the rate of multiple on-target insertions from 4% to 53% of clones (**Extended Data Fig. 9B**). Due to HEK293FT’s pseudo-triploid genome and copy number variation/instability ^33^, on-target insertions ranged from 0-5 per cell, with a median of 2 for both hifiDn29-dCas9 and Dn29-dCas9 clones (**Extended Data Fig. 9C**). Overall, these single cell results demonstrate hifiDn29-dCas9’s enhanced precision and efficiency, nominate hifiDn29 for generating clonal cell lines containing single on-target integrations, and highlight the value of single-cell analysis in evaluating gene editing outcome heterogeneity.

### Engineered LSR systems insert multi-kilobase DNA cargo into the genome of non-dividing cells, human embryonic stem cells, and primary T cells

Next, we sought to benchmark our engineered LSRs in a diverse set of genome insertion tasks across non-dividing cells and dividing primary cells. LSRs offer an advantage in genome engineering applications involving non-dividing cells due to their independence from DNA repair machinery and homologous recombination ^34^. We treated HEK293FT cells with aphidicolin to induce cell cycle arrest and observed that Dn29 and key variants showed largely equivalent integration rates to untreated cells. dCas9 fusions experienced decreased integration efficiency but still achieved up to 30% on-target integration (**Fig. 6A**).

**Figure 6:**
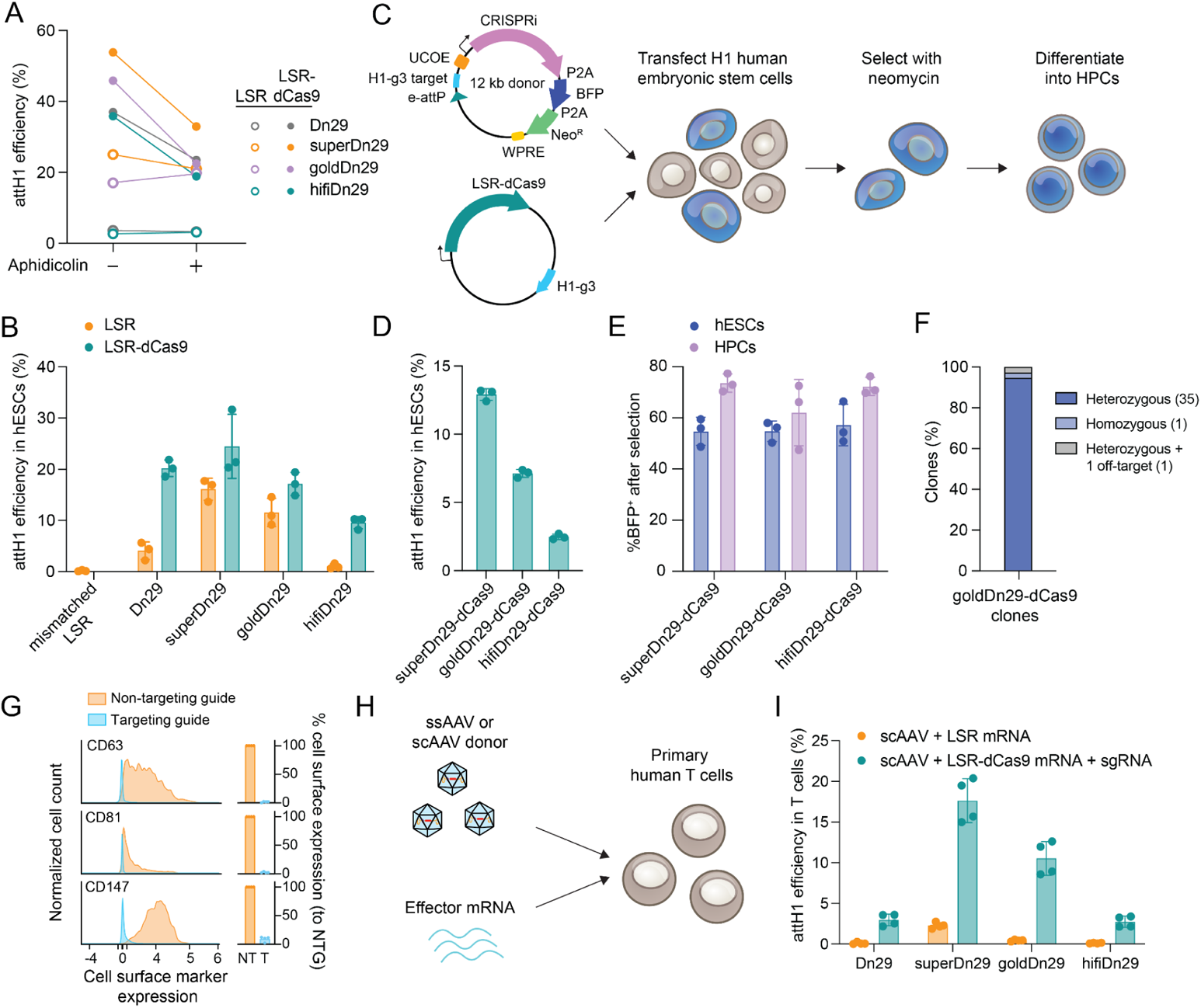
Engineered LSR systems insert large DNA cargo into the genome of non-dividing cells, human embryonic stem cells, and primary T cells. A. Integration efficiencies of Dn29 variants and dCas9 fusions at attH1, with and without cell cycle arrest by aphidicolin treatment. The dots represent the mean of n=3 biological replicates. B. Integration efficiencies of Dn29 variants and dCas9 fusions in human embryonic stem cells (hESCs). Bars and error bars represent the mean ± SD of n=3 biological replicates, shown as dots. C. Schematic of hESC engineering with LSRs. A 12 kb CRISPRi donor plasmid and LSR-dCas9 effector/guide plasmid is transfected into H1 hESCs, selected, and differentiated into hematopoietic progenitor cells (HPCs). UCOE: Ubiquitous Chromatin Opening Element, WPRE: Woodchuck Hepatitis Virus Posttranscriptional Regulatory Element. D. Integration efficiency of 12 kb CRISPRi donor by LSR-dCas9s at attH1. Bars and error bars represent the mean ± SD of n=3 biological replicates, shown as dots. E. CRISPRi-BFP cassette expression in engineered hESCs after selection, pre- and post-differentiation into HPCs. Bars and error bars represent the mean ± SD of n=3 biological replicates, shown as dots. F. Genotyping hESC single cell clones engineered with goldDn29-dCas9. N of clones analyzed per sample is indicated in the legend. G. HPC cell surface marker expression after guide transduction and selection, relative to non-targeting guide, in CRISPRi cells engineered by goldDn29-dCas9. Left histograms are representative cell surface marker expression plots. Right bar plots show the knockdown quantification of 4 biological replicates, calculated as target/non-target median fluorescence intensity, represented as a percentage. H. Schematic of ssAAV or scAAV donor and effector mRNA delivery into primary human T cells. I. Integration efficiencies of Dn29 variants and dCas9 fusions at attH1 in primary human T cells using scAAV donor. Bars and error bars represent the mean ± SD of n=4 biological replicates, each originating from a different blood donor.

We then tested DNA cargo installation in H1 human embryonic stem cells (hESCs) and observed that on-target integration efficiency with engineered recombinases increased up to 6-fold relative to wildtype Dn29, from 4.1% to 24.5% (**Fig. 6B**). These improvements are 25-50% of the integration efficiencies seen in HEK293FT, likely due to the 50-80% reduction in plasmid transfection efficiency in stem cells (**Extended Data Fig. 9D**). Insertion specificity also significantly improved, as off-target integration at attH3 for the engineered variants approached the ddPCR detection limit (**Extended Data Fig. 9E**).

To evaluate our optimized LSR variants’ capacity for larger cargo installation and enable functional genomics applications, we designed a large 12kb CRISPRi construct encoding for the dCas9-ZIM3 fusion and multiple regulatory elements and marker genes, including a BFP, neomycin resistance marker, Woodchuck post-transcriptional regulatory element (WPRE), and ubiquitous chromatin opening element (UCOE). We achieved robust insertion efficiencies up to 13% with standard lipid transfection (**Fig. 6C-D**).

After neomycin selection, ~60% of the bulk population was BFP^+^ by flow cytometry (**Fig. 6E**), indicating successful CRISPRi construct integration and expression. Clonal analysis of the goldDn29-dCas9-edited hESCs demonstrated that the majority (84%) of the clones were 100% BFP^+^, while some clones had significant levels of BFP silencing at the stem cell stage (**Extended Data Fig. 9F**). Genotyping revealed that 95% of clones possessed precise heterozygous attH1 insertions, with only one homozygous clone and one clone showing both on- and off-target integrations (**Fig. 6F**), These results underscore the high clonal consistency achieved, demonstrating a predominance of accurate, single-copy integrations and strong maintenance of cargo gene expression in H1 stem cells.

Transgene silencing is a persistent challenge in stem cell engineering, likely due to the extensive chromatin restructuring that occurs during stem cell differentiation ^35^. To assess whether attH1 supports stable cargo expression during differentiation, we differentiated the LSR-dCas9-edited hESCs into hematopoietic progenitor cells (HPCs) under neomycin selection. Post-differentiation, edited HPCs maintained robust cargo expression (~70% BFP+) with ~80% of cells expressing canonical HPC markers such as CD34 and CD43 (**Fig. 6E**, **Extended Data Fig. 10A**). Next, we introduced sgRNAs targeting CD63, CD81, and CD147 via lentiviral transduction into the goldDn29-dCas9-edited HPCs and observed 91-98% knockdown of these cell surface markers compared to a non-targeting guide (**Fig. 6G**, **Extended Data Fig. 10B**). Taken together, we demonstrate that engineered recombinases efficiently produce bulk hESC lines with near-clonal genotypes and stable large cargo expression throughout hESC-to-HPC differentiation, making them suitable for generating stable cell lines for CRISPR screens or potentially introducing therapeutic genetic cargoes.

Site-specific transgene insertions show great promise in immune cell engineering, enabling integration of functional cargos like chimeric antigen receptors (CARs) and additional immune regulators for therapeutic applications. However, high plasmid DNA toxicity in primary T cells (**Extended Data Fig. 10C**) limits use of conventional donor and effector expression plasmids. To address this, we electroporated T cells with effector mRNA and delivered donor templates via single-stranded AAV (ssAAV) or self-complementary AAV (scAAV) expressing mCherry (**Fig. 6H-I**, **Extended Data Fig. 10D-F**). LSR-dCas9 fusions achieved up to 17% integration efficiency into attH1 using AAV donors, which maintained high cell viability even at the highest doses. The scAAV donor yielded 2.3 to 3.3-fold higher integration rates compared to the ssAAV donor, likely due to the requirement of double-stranded DNA for LSR-mediated integration. The efficiency and viability we observe strongly support further development of this approach for engineering primary T cells.

Finally, we investigated the cross-reactivity of our engineered LSRs across model organisms, including mice and various non-human primates. An attH1-like sequence is present in the NEBL intron in marmosets, rhesus monkeys, and cynomolgus monkeys, with 1-2 point mutations compared to attH1, and is located intergenically in the mouse X chromosome with 6 point mutations. Dn29 and goldDn29 could robustly recombine attP with the model organism pseudosites in a plasmid recombination assay (**Extended Data Fig. 10G**), enabling future advancement in preclinical animal studies that bridge the gap between laboratory research and human clinical trials.

## Discussion

Here, we report a framework that combines directed evolution, protein engineering, and ML models for engineering DNA recombinases to efficiently and specifically insert large genetic cargos directly into the human genome, overcoming the need to preinstall attachment site landing pads. As a proof-of-concept, we report integration into a single genomic locus using optimized Dn29 LSRs, achieving a 13- to 17-fold improvement in insertion efficiency. Combining mutants with dCas9 fusions and optimized donor sequences (e-attP and sgRNA target sites) yielded recombinases with 40-53% efficiency and 90-97% genome-wide specificity for an endogenous locus.

Our engineering efforts provide numerous mechanistic insights into LSR function during genome integration. Notably, our dCas9 fusion experiments demonstrate that improved genome search and DNA binding are crucial areas for increased integration efficiency. To further interrogate DNA binding, we employed structural modeling and attachment site screening to identify specific protein and DNA regions critical for target recognition. Directed evolution revealed an inherent trade-off between efficiency- and specificity-improving mutations, which we overcame by strategically pairing mutations across distinct LSR domains and increasing the protein-DNA interface through fusions with DNA-binding proteins.

Our current system incorporates CRISPR components, which include both protein (dCas9) and RNA (sgRNA) elements. Although this design improves efficiency by 7-fold and specificity by 5-fold, it also increases the overall size of the system and introduces an additional RNA component, which increases the manufacturing and formulation complexity. In future iterations, these CRISPR components could be replaced with smaller, protein-only DNA binding domains such as zinc fingers. Such modifications would preserve the delivery advantages of these compact recombinases and maintain a streamlined system of a single protein and single DNA donor.

During the preparation of this manuscript, independent efforts to engineer the Bxb1 LSR were recently reported ^36,37^. These studies focused on improving Bxb1 targeting of its natural attB site to enhance integration rates into landing pads or retarget Bxb1 to endogenous sites. However, these approaches require pre-installation of the landing pad or concurrent delivery of multiple Bxb1 variants, increasing delivery complexity and increasing the space of potential off-target sites. In contrast, our study presents multiple orthogonal engineering strategies to enhance an LSR’s ability to recognize and integrate at endogenous genomic sequences. We demonstrate the generalizability of these approaches beyond Dn29 to Nm60, improving on-target genomic insertion efficiency to 73%.

These advancements have utility across diverse research and therapeutic applications of LSRs. The current paradigm for functional genomics employs lentiviral engineering of cell lines and single copy installation of pooled libraries, which can lead to unpredictable effects on gene expression, potential insertional mutagenesis, and silencing of transgenes^38^. Our engineered recombinases overcome these limitations by targeting a defined integration locus at high efficiencies, which are essential for large-scale and uniform functional genomics studies. In hESCs, we demonstrate that 95% of cells have single copy, on-target insertions, enabling the generation of homogenous bulk cell populations without single-clone selection. Furthermore, we previously demonstrated the utility of LSRs for virus-free library screening in landing pad cell lines^13^. Our new work extends this capability, showing the feasibility of integrating guide or protein libraries directly into an endogenous human genomic locus. In hESCs, these integrations occur at copy numbers comparable to low multiplicity of infection (MOI) lentivirus (MOI=0.1), well within the standard guidelines for approximating one integrant per cell^39^.

In the therapeutic space, our approach offers advantages over prevailing CRISPR-based gene therapies, which require a new guide RNA to target a distinct disease-causing mutation. By contrast, multi-kilobase insertions enable replacement of entire corrective open reading frames, providing a “one size fits all” approach for correcting genetic diseases with mutational heterogeneity across patient populations. Additionally, these corrective transgenes can include critical non-coding regulatory elements for enhanced control of gene expression. Furthermore, Dn29 can cross-reactively integrate into attH1-like sequences in diverse model organisms, an important consideration for future IND-enabling studies.

The strategies outlined in this work can be adapted to target diverse genomic loci beyond Dn29 attH1. To target a different locus, such as validated genomic safe harbors like *AAVS1* or therapeutic targets like *TRAC*, LSRs can be mined from the thousands of naturally occurring orthologs to find a recombinase with a closer match to the desired target sequence. These candidate LSRs can then be subjected to our joint optimization approach, combining directed evolution, machine learning predictions, and DNA-binding protein fusions to enhance both efficiency and specificity. Overall, the distinct LSR engineering strategies presented in this study provide a comprehensive blueprint for the development of next-generation recombinases capable of direct, site-specific genome integrations.

## Methods

### Ethics Statement

Our research complies with relevant ethical regulations. Experiments using human embryonic stem cell lines were performed under allowance granted by the Arc Institute Stem Cell Research Oversight Committee.

### Cell lines and culture

Experiments were conducted in HEK293FT cells (Thermo Fisher), H1 human embryonic stem cells (hESCs), and primary human T cells (STEMCELL Technologies) from deidentified healthy donors (Cat #200-0092). HEK293FT cells were cultured in DMEM with 10% FBS (Gibco) and 1x Penicillin-Streptomycin (Thermo Fisher), and dissociated using TrypLE Express (Gibco). H1 hESCs were maintained in MTeSR Plus (STEMCELL Technologies) supplemented with 1x Antibiotic-Antimycotic (Thermo Fisher) and cultured on Cultrex (Bio-Techne) or Matrigel (Corning) coated plates. For routine passaging, hESCs were dissociated with ReLeSR (STEMCELL Technologies). For 96-well plating prior to transfections, single-cell dissociation was performed using Accutase (STEMCELL Technologies). H1 hESCs were supplemented with 10 µM Rock inhibitor for 24 hours post-dissociation. Primary human T cells were cultured in complete X-VIVO 15 (cXVIVO 15) (Lonza Bioscience, Visp, Switzerland #04-418Q) which consists of 5% FCS (R&D systems, Cat #M19187), 5 ng μl^−1^ IL-7, and 5 ng μl^−1^ IL-15.

### Dn29 deep mutational scan library construction

An NNK deep mutational scanning library of the entire Dn29 CDS was generated using NNK oligos and overlap extension PCRs. First, forward and reverse oligos with NNK mixed bases at each codon were designed with a Tm of 65°C. Each NNK forward primer was paired with Dn29 DMS_universal_reverse that binds downstream of the CDS, and each NNK reverse primer with DMS_universal_forward primer, generating amplicons flanking the mutated codon. PCR reactions contained 2.5 µL Q5 Mastermix (NEB), 0.01 µL Dn29 plasmid template (100 ng/µL), 0.025 µL universal primer (100 µM), 1.465 µL water, and 1 µL unique NNK primer (2.5 µM). Cycling conditions: 98°C for 30 seconds; 30 cycles of 98°C for 10 seconds, 60°C for 30 seconds, and 72°C for 1 minute; final extension of 72°C for 2 minutes.

Upstream and downstream amplicons (2.5 µL each) were pooled and cleaned with 2 µL ExoSAP-IT (Thermo Fisher) and 0.5 µL of DpnI (NEB), incubating at 37°C for 30 minutes then 80°C for 15 minutes. For the overlap extension PCR, 1 µL cleaned PCR pool was mixed with 2.5 µL Q5 2x Mastermix, 0.025 µL of each universal primer (100 uM), and 1.45 µL of water, using the same cycling conditions.

The full mutant pool was created by combining 2.5 µL of each overlap extension PCR. The full-length Dn29 fragment was gel-extracted (Monarch DNA Gel Extraction Kit, NEB). The library and pEVO backbone were digested with XbaI and HindIII-HF (NEB). Ligation used 100 ng total DNA (3:1 molar ratio of library to backbone), 2 µL T4 ligase (NEB), 4 µL 10x T4 ligase buffer (NEB), and water to 40 µL. The reaction was split into two 20 µL reactions, ligated for 30 minutes at room temperature, inactivated at 65°C for 10 minutes, and purified (Clean and Concentrator-5 Kit, Zymo).

The ligation product was electroporated into XL-1 Blue cells (Agilent) according to the manufacturer’s instructions, recovered for 1 hour at 37°C in 1 mL SOC media, and plated onto four 245 mm × 245 mm Bioassay dishes. Approximately 1M colonies were obtained. Plasmids were purified using NucleoBond Xtra Midi EF kit (Macherey Nagel) and sequenced with Illumina NextSeq2000 600 cycle P1 kit (Figure S1C).

### Substrate-linked directed evolution

Library transformation, induction, and growth: 4 µL pEVO plasmid library was electroporated into 50 µL XL-1 Blue competent cells (Agilent), recovered in 1 mL SOC media (37°C, 1 hour), and then seeded into 100 mL LB media with carbenicillin and L-arabinose (10 µg/mL or 0 µg/mL). Cultures were grown overnight at 37°C. Library coverage (>1M colonies) was confirmed by plating serial dilutions. Plasmids were extracted using Qiagen Plasmid Midi kit (0.3 g wet bacteria pellet per column).

Selection of active variants: 500 ng plasmid was digested with NdeI (NEB) to eliminate inactive variants. Active variants were amplified using: 25 µL 2x Platinum Superfi II Mastermix (Thermo Fisher), 19 µL water, 2 µL each SLiDE_recovery_forward and SLiDE_recovery_reverse primers (10 µM), and 2 µL NdeI-digested material. PCR conditions: 98°C for 30 seconds; 30 cycles of 98°C for 10 seconds, 52°C for 10 seconds, 72°C for 55 seconds; final extension at 72°C for 5 minutes. The correct-size band was gel-extracted (Monarch DNA Gel Extraction Kit, NEB).

Cloning for next evolution cycle: Amplified active variants and pEVO backbone were digested with XbaI and HindIII-HF (NEB) at 37°C for 30 minutes, then heat-inactivated at 80°C for 20 minutes. Digested variants were purified using DNA Clean and Concentrator-5 (Zymo), and backbone with DNA Clean and Concentrator-25 (Zymo). Five ligation reactions (20 µL each) were set up using 100 ng DNA (3:1 ratio of library to backbone) and T4 ligase (NEB). Ligation occurred at room temperature for 30 minutes, followed by heat inactivation at 65°C for 10 minutes. Pooled reactions were purified (DNA Clean and Concentrator-5 Kit, Zymo), eluted in 6 µL water, and electroporated into XL-1 Blue cells to start the next evolution cycle.

### DNA shuffling and fragment reassembly

Shuffling the active variants between rounds of cycling involved a uridine exchange PCR to partially exchange thymidines for uridine, USER enzyme fragmentation at uridine sites, primerless PCR fragment reassembly, and PCR for full-length gene recovery.

Uridine exchange PCR: Fragment size and yield was optimized by modifying dUTP/dTTP ratio, with the optimal ratio being 3/7. PCR mixture: 5 µL 10x Thermopol Buffer, 1 µL 10mM dNTPs, 1 µL each SLiDE_recovery_forward and SLiDE_recovery_reverse primers (10 µM), 1 µL plasmid library, 1 µL Taq Polymerase, and 40 µL water. Cycling conditions: 95°C for 30 seconds; 30 cycles of 95°C for 20 seconds, 60°C for 30 seconds, 68°C for 1 min/kb; final extension at 68°C for 5 minutes. Full-length gene band was gel-extracted (Monarch Gel Extraction Kit, NEB). USER enzyme digestion: 500 ng aliquots were digested with 2 µL USER Enzyme (NEB) at 37°C for 3 hours. Gel electrophoresis confirmed fragment distribution (100-1000bp).

Fragment reassembly: Fragments were purified (DNA Clean and Concentrator-5, Zymo) and reassembled in a Primerless PCR reaction using the following conditions: 25 µL purified fragments, 25 µL 2x Q5 High Fidelity Master Mix (NEB). Cycling conditions: 98°C for 30 seconds; 30-50 cycles of 98°C for 10 seconds, 30°C for 30 seconds (+1°C/cycle), 72°C for 1 minute (+4s/cycle); final extension at 72°C for 10 minutes. A final PCR was performed to recover only the full length Dn29 CDS for further rounds of directed evolution. The following conditions were used for full-length gene recovery: PCR mixture: 25 µL Platinum Superfi II 2x Mastermix (Thermo), 10 µL reassembled fragments, 2 µL each DMS_universal_forward and DMS_universal_reverse primers (10 µM), and 11 µL water. Cycling conditions: 98°C for 30 seconds; 35 cycles of 98°C for 10 seconds, 60°C for 10 seconds, 72°C for 55 seconds; final extension at 72°C for 5 minutes.

The gel-extracted, shuffled, and reassembled genes were cloned into the plasmid backbone using XbaI and HindIII digest and T4 ligation as previously described.

### Variant library next generation sequencing and analysis

Six primer sets (DMS_NGS primers, see **Table S5**) were designed to amplify ~260 bp segments of the Dn29 CDS with Illumina adapter overhangs. Two rounds of PCR were performed to add P5/P7 adapters and i5/i7 indexes (FLAP2 primers). Amplicons were cleaned with Ampure XP beads (Beckman Coulter) between PCR rounds and after the final PCR. Amplicons were pooled in equimolar ratios, quantified using Qubit dsDNA High Sensitivity Kit (Thermo Fisher), and sequenced on Illumina NextSeq2000 (600 cycle kit). Full overlap between read 1 and read 2 was ensured for higher confidence in mutation calling.

Paired-end reads were merged using BBMerge (v.39.06) and analyzed with a custom Python script. The script converted Phred Quality scores to error probabilities using the formula *P* = 10^(*Q*/-10)^, where P is the probability of error and Q is the Phred Quality Score. Reads with a summed error probability greater than 0.5 or containing frameshifts were filtered out. Nucleotide and amino acid mutations at each position were then counted and plotted. Enrichment for each amino acid (AA) between the input and output libraries was calculated using the formula: ((%AA_output_)/(1-%AA_output_))/((%AA_input_)/(1-%AA_input_)). To distinguish library construction-based dropouts from selection-based dropouts in the enrichment heat maps, any amino acids with zero reads in the output library were assigned a single read.

### Nanopore sequencing and analysis

Variants were cloned into a vector containing a 100 nucleotide random UMI barcode with a BHVD repeat pattern. The plasmid library was linearized by Eco105I digestion. Nanopore libraries were prepared using the barcoded nanopore sequencing kit (SQK-NBD114.24) with 1 µg linearized plasmid library and sequenced on a MinION flow cell (R10.4.1) for 72 hours. Sequencing reads were filtered using nanoq (v.0.9.0) with settings *--min_len 4500 --max_len 5500 --min-qual 10* (i.e. minimum q score of 10, a minimum read length equivalent to 90% of the expected read length, and a maximum read length equivalent to 110% of the expected read length). The UMI sequence was extracted using cutadapt (v.1.18) with settings *–g “GGCGGTCACCATCACCACCACCACGCTACACG;max_error_rate=0.2…ACTGTAC;max_error_rate=0.2” --trimmed-only --revcomp --minimum_length 95*. All UMI sequences were trimmed to 95 nt using seqkit (v.1.3-r106) with the command *seqkit subseq -r 1:95*.

Reads were clustered by UMI with mmseqs easy-linclust (v.14.7e284) with setting *--min-seq-id 0.5*. For each UMI cluster bin with at least 15 reads, a representative cluster sequence was generated by using usearch (v.11) with settings *-cluster_fast -id 0.75 -strand both -sizeout -centroids* and taking the first representative sequence of the output ^40^. A final consensus sequence was generated by 1 round of polishing with Medaka (v.1.9.1) with settings *-m r1041_e82_260bps_hac_g632*. Counts for each unique variant were determined by tallying the total consensus sequences.

### Cloning variant library into a mammalian expression vector

Primers (DE_mammalian_forward and DE_mammalian_reverse) were designed to amplify the Dn29 CDS from the active variant PCR library, adding overhangs for Esp3i-compatible Golden Gate cloning. PCR conditions: 25 µL 2x Platinum Superfi II Mastermix (Thermo Fisher), 19 µL water, 2 µL purified active variant library, 2 µL each primer. Cycling: 98°C for 60 seconds; 30 cycles of 98°C for 10 seconds, 60°C for 10 seconds, 72°C for 55 seconds; final extension at 72°C for 5 minutes. The product was purified (DNA Clean and Concentrator-5, Zymo) and quantified by Nanodrop.

A mammalian expression vector was designed with the EF1α promoter upstream of an Esp3i golden gate landing pad, used as the destination for the protein variant library. The landing pad was followed by a T2A self-cleaving peptide sequence and an EGFP CDS.

Golden Gate reaction mixture: 75 ng mammalian expression vector, amplified variant library (3:1 molar ratio to vector), 1 µL T4 DNA Ligase Buffer (NEB), 0.5 µL T4 DNA Ligase (NEB), 0.5 µL Esp3i (ThermoFisher), and up to 10 µL nuclease-free water. Cycling: 35 cycles of 37°C for 1 minute, 16°C for 1 minute; 37°C for 30 minutes; 80°C for 20 minutes. Five Golden Gate reactions were performed, pooled, and purified (DNA Clean and Concentrator-5, Zymo). The library was transformed into Mach1 *E. coli* and plated for overnight growth. Random colonies were picked, grown in 4 mL TB-Carbenicillin, and miniprepped (NucleoSpin Plasmid Transfection Grade Mini kit, Machery-Nagel).

### Transfection of HEK293FTs for assessing genomic integration

One day before transfection, 12-18K HEK293FT cells were plated per well of a 96-well plate, aiming for 60%-80% confluency at the time of transfection.

Standard LSR + donor transfection: for transfections containing an LSR effector plasmid and a donor plasmid, each well was transfected with 725 ng of DNA, containing a 5:1 molar ratio of donor plasmid to effector plasmid, using 0.5 µL of Lipofectamine 2000 (Thermo) per well.

Standard LSR-dCas9 + donor + guide transfection: LSR-dCas9 effector plasmid, donor plasmid, and guide plasmid were transfected with 725 ng total DNA, containing a 5:1:1 molar ratio of donor:effector:guide plasmid with 0.5 µL of Lipofectamine 2000 per well, unless specified otherwise in figure legends.

Modified transfection conditions: Experiments shown in **Fig. 3D, 3E, and S7B** were transfected with 375 ng of effector plasmid, 100ng sgRNA plasmid, and 250 ng donor plasmid using Lipofectamine 2000. Experiments shown in **Fig. 3G, 5 and 6** used a consolidated plasmid expressing both the effector and guide RNA. In HEK293FT experiments, this consolidated plasmid was transfected at a 5:1 ratio of donor:effector/guide plasmid with 0.585 µL Lipofectamine 2000 per well. For transfections containing two gRNA plasmids (**Fig. 3K, S8B**), each well was transfected with 375 ng effector plasmid, 75 ng each gRNA plasmid, and 250 ng donor plasmid.

The cells were incubated and monitored for three days for mCherry (donor plasmid) and GFP (effector plasmid) expression. Cells were then harvested for flow cytometry (Attune NxT Flow Cytometer, Thermo Fisher) or genomic DNA extraction for downstream analyses.

### Cell Harvest, ddPCR, qPCR and flow cytometry

Three days post-transfection, cells were trypsinized with 50 µL TrypLE (Gibco) for 10 minutes and then quenched with 50 µL Stain Buffer (BD). The 100 µL cell suspension was split into two 50 µL aliquots in U-bottom 96-well plates, centrifuged (300 × g, 5 minutes), and supernatant aspirated. One plate was resuspended in 200 µL Stain Buffer (BD) and analyzed with Attune NxT Flow Cytometer with Autosampler (Thermo Fisher).

The other plate was resuspended in 50 µL QuickExtract DNA Solution (Biosearch Technologies), vortexed for 15 seconds, and thermocycled: 65°C for 15 minutes, 68°C for 15 minutes, 98°C for 10 minutes. DNA was cleaned with 0.9x AmpureXP (Beckman Coulter) beads.

To assess integration efficiency and specificity, qPCR/ddPCR primers and probes were designed to span the left integration junction of attH1 and attH3, using a constant primer that binds to the donor plasmid sequence (ddPCR_donor_reverse_1) a genome binding primer near the pseudosite (ddPCR_attH1_forward_1, ddPCR_attH3_forward) and a FAM probe within the amplicon (ddPCR_attH1_probe_1, ddPCR_attH3_probe). For attH1, a second set of primers/probes was designed to target the right junction to verify measurement accuracy (ddPCR_attH1_2 primers/probe). Genomic reference primers and probes located nearby each attachment site were designed to measure pseudosite copy number for efficiency percentage calculations.

ddPCR reaction mix (22 µL total): 11 µL ddPCR Supermix for Probes (no dUTP) (Bio-Rad), 1.98 µL of each primer (10 µM), 0.55 µL of each probe (10 µM), 1.65 µL cleaned gDNA, 0.22 µL SacI-HF (NEB), water to volume. Each reaction contained primers and probes for the target site (FAM probe) and a nearby reference locus (HEX probe). Reactions were run on QX200 AutoDG Droplet Digital PCR System (Biorad). For off-target detection or low concentration samples, primers were increased to 20 µM and volume halved, and gDNA volume was increased to 4.95 µL.

qPCR reaction mix (40 µL total): 1 µL of each primer, 0.8 µL of each probe, 20 µL TaqMan Fast Advanced Mastermix (Thermo Fisher), 2.4 µL genomic DNA, and 12 µL water. Mastermix was split into three 10 µL technical replicates in a 384-well plate and run on LightCycler 480 (Roche). Primer pairs for ddPCR and qPCR are provided in **Table S5**.

### Three plasmid recombination assay in HEK293FT cells

A fluorescent reporter assay was used to assess episomal plasmid recombination in HEK293FT cells. One day before transfection, 12-18K HEK293FT cells were plated per well of a 96-well plate, aiming for 60%-80% confluency at the time of transfection. Three plasmids at a 1:1:1 molar ratio were transfected into the cells using Lipofectamine 2000: 1. 200 ng of the effector plasmid expressing the Dn29 variants and GFP, 2. 50.5 ng of the donor plasmid containing the attP attachment sequence and mCherry, and 3. 70.6 ng of the acceptor plasmid containing an Ef1a promoter and the cognate attB attachment sequence. Upon recombination of the two attachment sequences, the Ef1a promoter will drive expression of the mCherry CDS, which is read out by flow cytometry (**Fig. S6D**). To assess the excision reaction, the attP in the donor plasmid is replaced with the left post-recombination attachment site (attB-L:attP-R), called attL, and the attB is replaced with the right post-recombination attachment site (attP-L:attB-R), called attR. To assess attP recombination with model organism pseudosites, the attB sequence is replaced with the pseudosite sequences. Mismatching LSR (Bxb1) controls with each donor and acceptor plasmid is used to correct for the leaky mCherry background expression, defining the flow cytometry gating boundaries. Three days post-transfection, the cells were trypsinized with 50 µL TrypLE (Gibco) for 10 minutes, quenched with 50 µL Stain Buffer (BD), transferred to U-bottom 96-well plates, centrifuged (300 × g, 5 minutes), and supernatant aspirated. Plates were resuspended in 200 µL Stain Buffer (BD) and analyzed with Attune NxT Flow Cytometer with Autosampler (Thermo Fisher).

### Site-directed mutagenesis for combinatorial mutant cloning

Site-directed mutagenesis (SDM) primers were designed using the script from Bi et al. 2020, selecting primers with Tm closest to 65°C ^41^. For each mutation, a forward and reverse primer were generated, each containing the desired mutation at the center. PCR reactions were set up combining: forward SDM primer with DMS_universal_reverse primer or reverse SDM primer with DMS_universal_reverse primer. PCR mixture (12.5 µL total): 6.25 µL Platinum Superfi II Mastermix, 0.5 µL each primer (10 µM), 1 µL plasmid template DNA (1 ng/µL), water to volume. PCR was run using the standard Platinum Superfi II Mastermix protocol with annealing temperature at 65°C. Products were cleaned with 0.5x Ampure XP beads. For Gibson assembly, 1 µL each cleaned PCR product, 5 µL Gibson mastermix, and 3 µL water was incubated at 50°C for 15 minutes, then transformed into Mach1 E. coli and plated. For simultaneous cloning of two or more mutations, universal primers were replaced with other mutation’s forward and reverse primers. Two mutations required a two-piece Gibson assembly, three mutations required a three-piece assembly, and so forth.

### Genome-wide integration site mapping

HEK293FTs were transfected as previously described, with a non-matching LSR (Bxb1) plasmid replacing the effector plasmid as a control for donor plasmid dilution. Cells were cultured for 2-3 weeks, passaging and analyzing by flow cytometry every 2-3 days at 80% confluency, until the non-matching LSR control was <1% mCherry+, indicating the plasmid had nearly completely diluted out. Genomic DNA was extracted using Quick-DNA Miniprep Plus Kit (Zymo), quantified by Qubit HS dsDNA Assay (Thermo), and 1 µg of gDNA per sample was DpnI-digested (NEB) to remove residual donor plasmid. Tn5 tagmentation, nested PCR enrichment of integration sites, NGS sequencing, and computational analysis were performed as described in Durrant et al. 2023. To reduce occurrence of index hopping, unique dual i7 and i5 barcodes were utilized for the attH1 targeted samples in Figure 3. To directly compare specificity of samples with different numbers of measured integration events, samples were downsampled to the same total UMI count.

### LSR-dCas9 and gRNA plasmid design and cloning

Fusion proteins consisting of a catalytically dead Cas9 fused to an LSR and a P2A-GFP were constructed by Gibson assembly into a pUC19-derived plasmid containing the Ef1a promoter and a SV40 poly-A tail. Variable flexible linkers, including a (GGS)_8_, (GGGGS)_6_ XTEN16, XTEN32-(GGSS)_2_, and XTEN48-(GGSS)_2_ were used to link the dCas9 and LSR. Spacers targeting loci proximal to the LSR integration site and non-targeting controls were cloned into an sgRNA expressing plasmid via oligo ligation and Golden Gate cloning. Spacer selection was based on PAM sequence and pseudosite proximity.

### Designing and cloning the attP library

Two plasmid libraries (attP-L and attP-R) were constructed to determine nucleotide preference within the attP, with each 26 bp half-site mutagenized separately. IDT-synthesized oligo pools contained 79% WT base and 7% each of the other bases at each position. Single stranded oligo pools were subjected to second strand synthesis. First, an oligo anneal reaction containing 2 µL Library Oligo (100 µM), 4 µL klenow primer (100 µM), 3.4 µL 10x STE Buffer, and 24.6 µL water was heated at 95°C for 5 minutes, then cooled to room temperature. Next, a Klenow extension reaction containing 34 µL annealed libraries, 8 µL water, 5 µL 10x NEBuffer2, 2 µL 10 mM dNTPs (NEB), and 1 µL DNA Polymerase I, Large (Klenow) Fragment (NEB, 5000 U/mL) was incubated at 37°C for 30 minutes, purified (DNA Clean and Concentrator-5, Zymo), and eluted in 20 µL nuclease-free water.

The purified product was cloned by Esp3i Golden Gate cloning into pCB235: 75 ng pre-digested backbone, 3:1 molar ratio of attP library to backbone, 0.5 µL each of T4 DNA ligase (NEB) and Esp3i (Thermo Fisher), 1 µL T4 DNA Ligase Buffer (NEB), and water to 10 µL was incubated at 37°C for 1 hour, purified, and eluted in 6 µL nuclease-free water. 1 µL purified library was electroporated into Endura Electrocompetent Cells (Biosearch Technologies) at 10 μF, 600 Ω, 1800 V, recovered in 2 mL Lucigen Recovery Media (37°C, 1 hour), plated on 245 mm × 245 mm Bioassay dishes, and incubated at 30°C overnight. Final libraries were scraped, purified (Nucleobond Xtra Maxi EF kit, Machery-Nagel), and sequenced with Illumina NextSeq2000.

### attP library transfection, harvest, and library preparation

2.2 × 10^6^ HEK293FT cells were plated on 10 cm dishes one day before transfection to achieve 70% confluence at transfection. 24 μg total plasmid DNA was prepared at a 5:1:1 molar ratio (attP library:LSR effector:sgRNA). DNA and 72 μL Lipofectamine 2000 were separately mixed with 1.5 mL OMEM, incubated for 5 minutes, then combined and incubated for 10 minutes before adding dropwise to cells. After three days, cells were harvested with TrypLE (Gibco) and genomic DNA extracted using Quick DNA Midiprep Plus Kit (Zymo).

Integration events were amplified by single-step PCR with i5/i7 index-adding primers using all available genomic DNA. Biological replicates had 1 bp staggered amplicons to increase nucleotide diversity. PCR conditions: 25 μL NEBNext High Fidelity PCR Master Mix, 2.5 μg genomic DNA, 1.25 μL each of the attL or attR i5 or i7 primers (**Table S5**), water to 50 μL. Cycling: 25 cycles of 98°C for 10s, 63°C for 10s, 72°C for 25s. PCR products were pooled, run on 2% agarose gel, and correct-size bands were extracted (Monarch® DNA Gel Extraction Kit, NEB). Libraries were quantified (Qubit 1x dsDNA High Sensitivity assay, Thermo Fisher), pooled equimolar with 35% PhiX spike-in, and sequenced on Illumina NextSeq2000 (150bp paired-end reads).

### attP Library enrichment analysis

Libraries were demultiplexed using Illumina Basespace automatic demultiplexing workflow. Paired-end reads were merged using BBMerge (v.39.06) and analyzed with a custom Python script. Reads were filtered for exact amplicon length and QScore ≥ 30. Next, percent abundance of each nucleotide at each attP position was calculated for input and output libraries. Enrichment scores were computed using the equation: 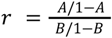, where A and B represent the read counts for selected nucleotides in output and input libraries, respectively, normalized to the total number of reads. Enrichment scores were converted to sequence logos, generated using Logomaker ^42^ and matplotlib packages.

Unique library members recovered as integration events were assessed by generating the set of unique reads. The number of unique integration events from NGS analysis was compared to ddPCR analysis of bulk genomic DNA for validation.

### Stem cell transfection

H1 hESCs were cultured in MTeSR Plus medium (STEMCELL Technologies) on Cultrex-coated (Bio-Techne) or Matrigel-coated (Corning) 6-well plates. Cells were routinely subcultured at a 1:12 ratio using ReLeSR Passaging Reagent (STEMCELL Technologies) every four days or at 70-80% confluency. Three days after splitting (60% confluency), the cells were dissociated for 10 minutes with Accutase (STEMCELL Technologies), and plated in Cultrex-coated 96-well plates at 25-30K cells per well with 10 µM Rock inhibitor. The next day (at 70% confluency), media was changed to include 50 µM Rock inhibitor two hours pre-transfection. 3 µg plasmid DNA containing a 1:1 molar ratio of combined effector/guide plasmid to donor plasmid in 10 µL volume was diluted in 81 µL MTeSR Plus and thoroughly pipette mixed. 9 µL of Fugene HD Transfection Reagent (Promega) was added to the DNA/MTeSR mix, thoroughly mixed, and incubated for 12 minutes. After another thorough pipette mix, 7 µL of the DNA was added dropwise to each well. The cells were incubated at 37°C for 1-2 days, splitting 1:2 if 90% confluency was reached. After 3 days, the cells were dissociated with Accutase and split into two V-bottom plates, one for flow cytometry and one for gDNA harvest with QuickExtract DNA Solution (BioSearch Technologies).

### HPC differentiation and surface marker staining

hESCs were differentiated into hematopoietic progenitor cells using the STEMdiff Hematopoietic Kit (STEMCELL Technologies). On day 10 of differentiation, 250 µL of non-adherent cells were collected from the supernatant using wide bore P1000 tips and transferred to a V-bottom 96 well plate. Next, the cells were pelleted at 400g for 5 minutes, supernatant discarded, and resuspended in 95 mL Stain Buffer (BD) containing 1 µL of each antibody with a wide bore pipette. The following antibodies were used: APC CD81 (BD, Cat:551112), APC CD147 (Thermo Fisher, Ref: A15706), Alexa Fluor® 647 CD63 (BD, Cat: 561983), APC/Cyanine7 CD34 (BioLegend, Cat: 343514), PE CD43 (BioLegend, Cat: 343204). The cells were incubated in the dark for 20 minutes to 1 hour, washed once with Stain Buffer, and flowed on the Attune Flow Cytometer (Thermo Fisher).

### hESC single cell dilution and genotyping

hESCs were diluted to 1 cell/100 µL in MTeSR Plus media supplemented with 1x CloneR (STEMCELL Technologies) and plated into two 96-well plates per sample. Cells were maintained until colonies were visible, then wells with multiple colonies were removed. Single colonies were expanded to 24-well dishes when they covered half the surface area of the 96-well. At the next split, one quarter of each well was pelleted for gDNA extraction using QuickExtract DNA Solution (BioSearch Technologies). The extracted gDNA was cleaned with 0.9x AmpureXP beads and genotyped by ddPCR. Primers and probes were designed to target the attH1 junction (ddPCR_attH1_1 set), the donor sequence (Amp_forward, Amp_reverse, Amp_probe), and a nearby genomic reference sequence. On-target zygosity was determined by the attH1/reference ratio, while total zygosity was measured by the donor/reference ratio.

### HEK293FT single cell sorting and genotyping

HEK293FT cells were transfected as previously described. Eight days post-transfection, cells were placed under puromycin selection (0.5 μg/mL) for 10 days. On day 18, cells were trypsinized, strained through a 35 µm filter, and single mCherry+ cells were sorted into four 96-well plates per sample using the FACSAria Fusion (BD). Single cell colonies were expanded for two weeks until >50% confluent, with visual inspection to ensure single colony growth. Wells with zero or multiple colonies were excluded from analysis.

Confluent colonies were harvested with QuickExtract DNA Solution (BioSearch Technologies) and amplified in two separate PCRs: PCR 1 using primers UMI_reverse and ddPCR_attH1_forward_1, flanking the UMI and attH1 donor/genome junction, and PCR 2 using primers UMI_reverse and UMI_forward, flanking the UMI on the donor plasmid. Amplicons were sequenced via Sanger and/or NGS to determine on-target UMI count (PCR 1) and total UMI count (PCR 2), allowing calculation of on-target and off-target insertion counts per colony.

### Lentivirus production and HPC transduction

sgRNA spacers targeting cell surface markers CD81, CD147, and CD63 were cloned into the LentiGuide-Puro construct (Addgene #52963). Lentivirus was generated using the LV-MAX Lentiviral Production Kit (Invitrogen) according to manufacturer’s instructions and concentrated 100x with Lenti-X Concentrator (Takara). HPCs were diluted to 50,000 cells per well in 100 µL of Media B (STEMdiff Hematopoietic Kit, STEMCELL Technologies) in a 96-well plate. Each well received 1 µL of LentiBOOST (SIRION Biotech) and 1 µL of lentivirus. Media was changed the following day. Four days post-transduction, a subset of cells was stained for cell surface markers. Remaining cells were treated with 1 μg/mL puromycin for four days to select for transduced cells, followed by cell surface marker staining.

### Generating Alphafold3 models of Dn29 bound to attB

The full-length wildtype Dn29 protein sequence and minimal attB-L or attB-R sequence (attB-L: GTAGACAAGGAAGGTAATGA; attB-R: GAAATAAGTTTGATAGATAT) were input into the Alphafold3 web server with the seed set to “auto”. Five models were generated for each query of Dn29 bound to an attB half-site. Outputs were manually inspected to ensure correct orientation of Dn29 bound to the half-site, with the dinucleotide core of the DNA proximal to the NTD. One model (Dn29 × attB-R) out of the 10 generated models met this criterion and was selected for further analysis. The chosen model was compared to the Listeria Integrase crystal structure of the LSR CTD and attP complex (pdb:4KIS). Despite 4KIS being bound to attP instead of attB, domain-wise comparisons showed strong alignment: RMSDs were 1.341 and 1.707 for the zinc-ribbon domain, and recombinase domain, respectively (**Fig. S6B**). Protein/DNA interface residues were identified with the InterfaceResidues pymol script using default settings.

### Predicting combinatorial mutations and feature importance with machine learning

The efficiency and specificity data of all Dn29 variants were split into a training and test set based on what round of experimentation they were generated in. The training set, called round 1, contained all variants from the two single mutation validation experiments, where mutations were tested individually on top of variant 127 (**Fig. 1G, Fig. S4C-E**) or variant 381 (**Fig. S4F**). The testing set contained all higher order combinations from the iterative rounds of driver mutation stacking (rounds 2-5). The efficiency (percent of integrations at attH1) was normalized to wildtype and specificity (ratio of attH1/attH3 activity) was log-transformed. The full amino acid sequences of the protein variants were one-hot encoded, and activity in the training set was modeled using linear regression, ridge regression, XGBoost, and CatBoost with the scikit-learn, xgboost, and catboost Python libraries.

For the ridge regression, optimal alpha was identified through minimization of the testing set R^2^ (alpha=0.8 for efficiency model, alpha=1.3 for specificity model). Hyperparameter optimizations were conducted for XGBoost and CatBoost by performing a randomized search, evaluating on negative mean squared error, using the following parameters: XGBoost: ‘n_estimators’: [100, 500, 1000], ‘learning_rate’: [0.01, 0.05, 0.1], ‘max_depth’: [3, 5, 7], ‘subsample’: [0.5, 0.6, 0.7, 0.8, 1.0], ‘colsample_bytree’: [0.7, 0.8, 1.0]; CatBoost: ‘iterations’: [100, 200, 500, 1000], ‘learning_rate’: [0.01, 0.05, 0.1, 0.2], ‘depth’: [4, 6, 8, 10], ‘l2_leaf_reg’: [1, 3, 5, 7, 9], ‘bagging_temperature’: [0, 1, 2, 3], ‘random_strength’: [1, 1.5, 2, 3], ‘border_count’: [32, 64, 128], ‘grow_policy’: [‘SymmetricTree’, ‘Depthwise’, ‘Lossguide’].

The following parameters were chosen for each model: XGBoost, specificity: ‘subsample’= 0.5, ‘n_estimators’= 1000, ‘max_depth’= 7, ‘learning_rate’= 0.1, ‘colsample_bytree’= 0.8; XGBoost, efficiency: ‘subsample’= 0.7, ‘n_estimators’= 100, ‘max_depth’= 7, ‘learning_rate’= 0.05, ‘colsample_bytree’= 0.7; CatBoost, specificity: ‘random_strength’= 1.5, ‘learning_rate’= 0.1, ‘l2_leaf_reg’= 1, ‘iterations’= 1000, ‘grow_policy’= ‘Depthwise’, ‘depth’= 4, ‘border_count’= 128, ‘bagging_temperature’= 1; CatBoost; efficiency: ‘random_strength’= 1.5, ‘learning_rate’= 0.1, ‘l2_leaf_reg’= 7, ‘iterations’= 500, ‘grow_policy’= ‘Lossguide’, ‘depth’= 4, ‘border_count’= 32, ‘bagging_temperature’= 2.

### In vitro transcription and purification of mRNA

Effector constructs were cloned into an in vitro transcription (IVT) plasmid as previously described ^43^. This plasmid contained a mutated T7 promoter, 5’ UTR, P2A EGFP, and 3’ UTR followed by a 145 bp polyA sequence. IVT templates were generated by PCR using primers oGX006 and oLGR009, which incorporate a polyA tail and correct the T7 promoter mutation. PCR reactions were performed using KAPA-HiFi HotStart 2x (Roche) master mix with 6.25 ng of plasmid template per 25 µL reaction. The PCR protocol involved annealing at 63°C, extending for 45 seconds per kb, and running for 18 cycles. The reactions were purified using 0.8x volume of SPRI beads and eluted into water. The purified PCRs were analyzed by gel electrophoresis and nanodrop to ensure correct size and determine concentration.

The IVT reactions were set up using the HiScribe T7 High-Yield RNA Synthesis Kit (New England Biolabs, Cat #E2040S), modified with full pseudo-UTP substitution using N1-Methyl-Pseudo-U (TriLink Biotechnologies, Cat #N-1081) and co-transcriptionally capped with CleanCap AG (TriLink Biotechnologies, Cat #N-7113). Each IVT reaction contained 5 mM ATP, CTP, GTP, and pseudo-UTP, 4 mM CleanCAP AG, 1x Transcription Buffer, 3.75 ng/µL DNA template, 1 U/µL Murine RNAse Inhibitor (New England Biolabs, Cat #M0314L), 0.002 U/µL yeast inorganic pyrophosphatase (NEB. Cat # M2403L), and 5 U/µL T7 RNA polymerase. Reactions were incubated for 2.5 hours at 37°C.

Next, the mRNA was purified using lithium chloride. To each reaction, 1.5X water and 1.25X 7.5M LiCl was added. The solution was chilled at −20°C for 30 minutes and then spun at max speed (16,000xg) for 15 minutes at 4°C. The supernatant was discarded, and the pellet was rinsed with 70% ice cold ethanol to remove residual salts. After another max speed spin for 10 minutes at 4°C, the mRNA was resuspended in water and stored at −80°C. The mRNA was analyzed on the Agilent Tapestation and by Qubit RNA High Sensitivity (Thermo) to ensure correct size and determine concentration.

### Electroporation of primary human T cells

Two days before electroporation, T cells were seeded at 1 × 10^6^ fresh cells/mL and activated with a 1:1 bead-to-cell ratio with anti-CD3/CD28 Dynabeads (Life Technologies, #40203D). On the day of electroporation, the beads were magnetically removed and the T cells were electroporated with 2 µg LSR-dCas9-P2A-EGFP mRNA and 2 µg sgRNA (Synthego) for LSR-dCas9 samples or 1 µg LSR-P2A-EGFP mRNA for LSR samples using the Lonza P3 Primary Cell Kit. Each electroporation contained between 0.5 × 10^6^ − 1 × 10^6^ cells in 20 µL total volume, and was electroporated using the 4D Nucleofector system and the DS-137 pulse code. Immediately after electroporation, 80 µL pre-warmed culture media was added to the Nucleocuvette strip, which was then incubated at 37°C for 15-30 minutes. Next, 2 × 10^5^ cells per condition were split into 96 well U-bottom plates in 100 µL of serum free media (TheraPEAK® X-VIVO®-15 Serum-free Hematopoietic Cell Medium, Cat #BEBP04-744Q) supplemented with 5 ng μl^−1^ IL-7, and 5 ng μl^−1^ IL-15. Cells were then transduced at an MOI of 1 × 10^5^ genome copies/cell with ssAAV or scAAV vectors of serotype 6 (AAV6) containing the e-attP sequence, attH1 sgRNA target sequence, and an mCherry expression cassette which were ordered from VectorBuilder. The next morning, cells were spun down at 300 × g for 5 minutes, the serum free media was removed, and cells were resuspended in 200 µl of fresh cX-VIVO. Cells were maintained and passaged as needed by the addition of cX-VIVO every 2-3 days.

### T cell staining, flow cytometry, and genomic harvesting

Three days post-electroporation, 50 µL of T cells were collected for staining and flow cytometry. Briefly, cells were centrifuged, washed once with 200 μl cell staining buffer, and stained with Ghost Dye™ Red 780 at a 1:1000 dilution (Tonbo, Cat #13-0865-T500) for 20 minutes in the dark at 4°C. The cells were measured using an Attune NxT Cytometer with a 96-well autosampler (Invitrogen) and analyzed using FlowJo for viability, mCherry fluorescence (expressed on the AAV) and GFP fluorescence (effector expression). The remaining 150 µL of T cells in culture were centrifuged at 300 × g for 5 minutes and the gDNA was harvested using QuickExtract DNA Solution (BioSearch Technologies) and analyzed by ddPCR as described above.

### Generative AI

AI language models (ChatGPT and Claude) were utilized for generating custom python scripts for data analysis and visualization, assistance with copy-editing, and infilling preliminary drafts of some sections based on an author-provided outline. All AI-generated content was thoroughly reviewed, edited, and verified by the authors.

## Extended Data Figures

**Extended Data Figure 1:**
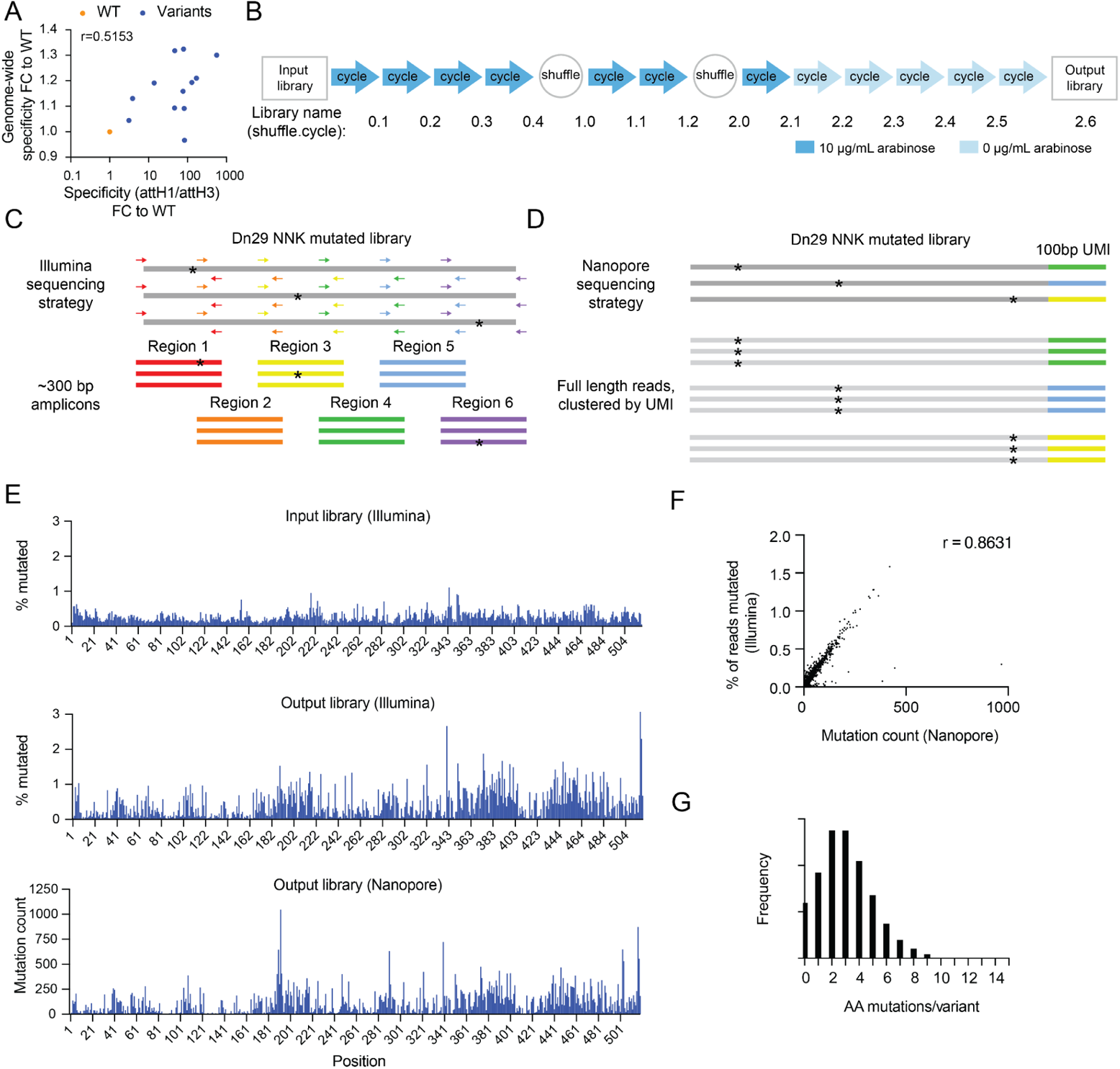
High-throughput sequencing and validation of directed evolution library. A. Correlation between genome-wide specificity (ratio of on-target insertions relative to all on- or off-target integration events) and specificity proxy (on-target/single off-target), shown as fold change to WT. Dots represent the mean of n=2 biological replicates. Pearson r=0.5153, one-tailed p=0.0297. B. Overview of selection cycles and DNA shuffling steps in directed evolution campaign. C. Illumina sequencing strategy for Dn29 NNK library. The 1557 bp CDS is sequenced in PCR fragments, generating 6 regions <300 bp each. D. Nanopore sequencing strategy for Dn29 NNK library. Library is cloned into N x 100 bp UMI plasmid backbone, and reads are clustered by UMI for consensus sequence generation. E. Percentage of mutated residues at each Dn29 coding sequence position: input library (Illumina) vs output library (Illumina and Nanopore). F. Correlation of mutational frequency in output library: Illumina vs Nanopore. Each dot represents a coding sequence position. Pearson r=0.8631, one-tailed p<0.0001. G. Distribution of amino acid mutations per variant in the output library (Nanopore).

**Extended Data Figure 2:**
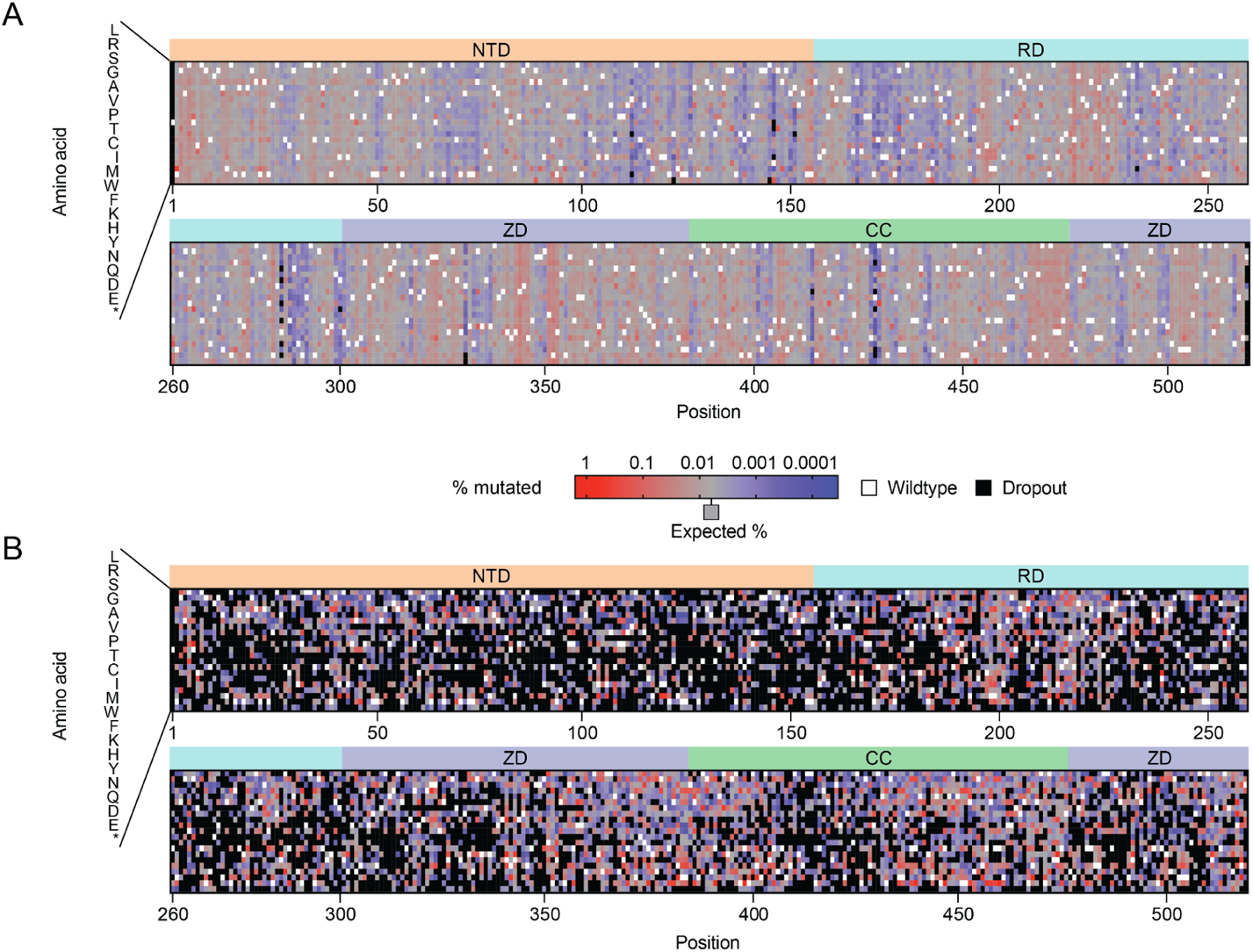
Mutation distribution of (A) input and (B) output libraries. White indicates the WT amino acid, black indicates an amino acid dropout, and red and blue indicate enrichment and depletion, respectively, compared to the expected mutation frequency (0.006%, shown in gray). Each value is normalized by the number of encoding codons in the NNK library. NTD: N-terminal domain; RD: recombinase domain; ZD: zinc-ribbon domain; CC: Coiled-coil domain.

**Extended Data Figure 3:**
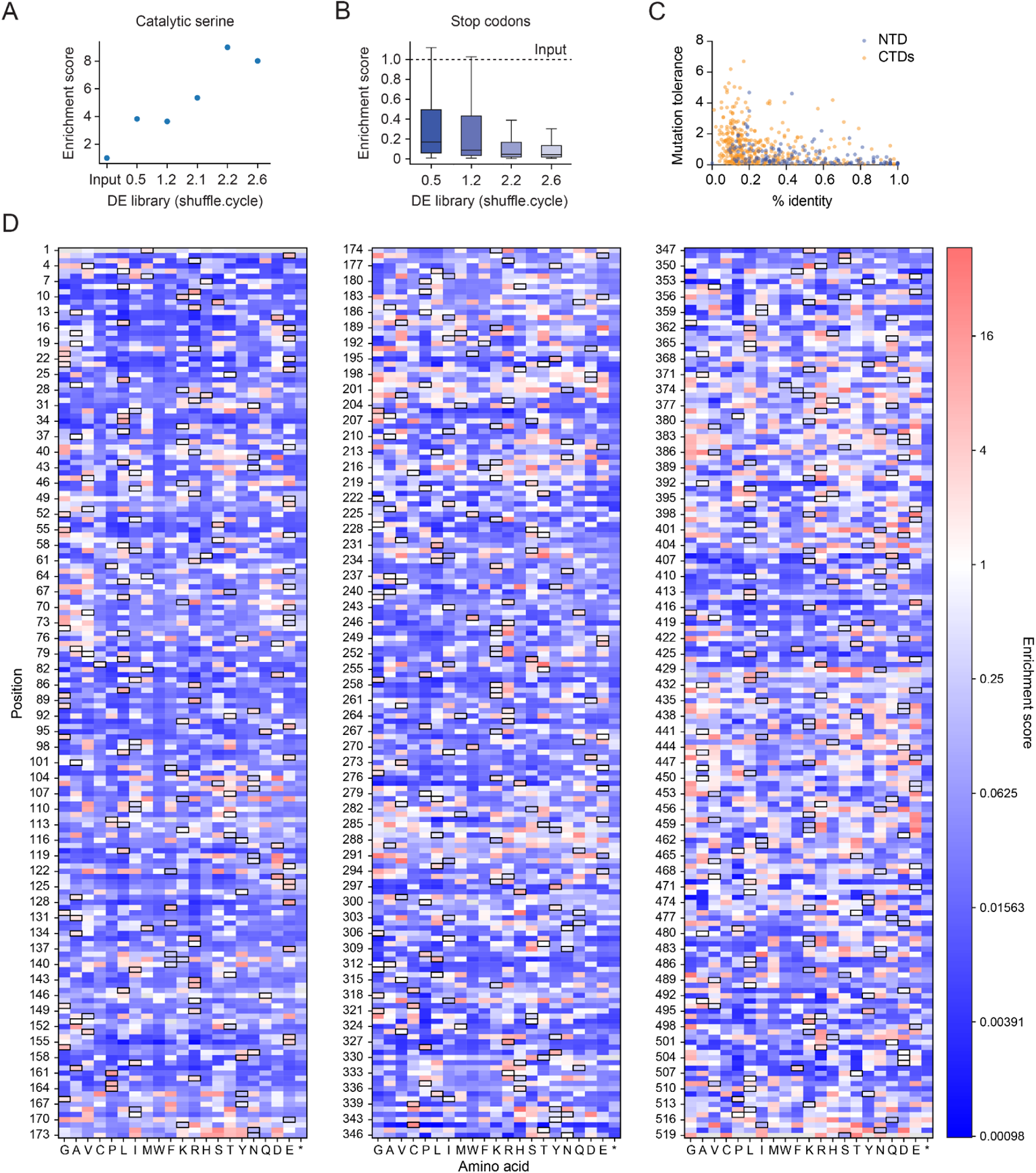
Mutational enrichment between input and output libraries. A. Enrichment score of the catalytic serine (10S) throughout various timepoints in the directed evolution campaign. B. Box plot of stop codon enrichment scores across all coding sequence positions at various time points. Boxes: interquartile range (IQR); line: median; whiskers: values within 1.5 times IQR. C. Correlation between mutational tolerance (average non-WT residue enrichment) and phylogenetic conservation (% identity from multiple sequence alignment of 106 LSR clusters within 30% identity of Dn29). Each dot represents a position in the CDS, colored by domain. NTD: N-terminal domain, CTDs: C-terminal domains. Pearson r = −0.3577, two-tailed p<0.0001. D. Enrichment score of output library normalized to input library. Boxed cells indicate WT amino acids, red and blue cells indicate enrichment or depletion, respectively, white cells indicate enrichment score of 1 (no enrichment), and gray cells represent an amino acid dropout in the input library.

**Extended Data Figure 4:**
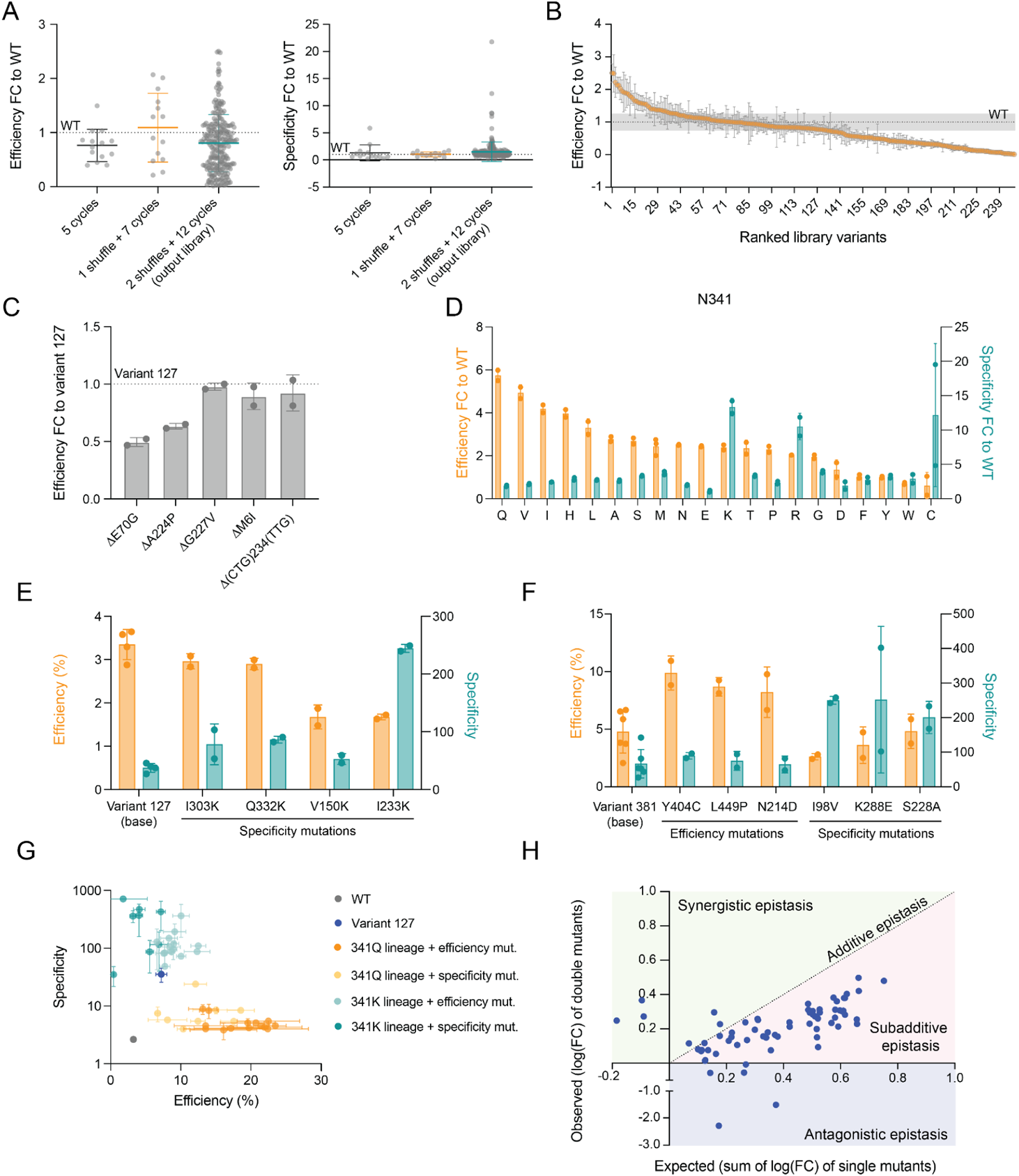
Exploration of mutational landscape with combinatorial mutations. A. Integration efficiency (left) and specificity (right) of variants from different libraries throughout directed evolution, as fold change to WT Dn29. Each dot represents the mean of n=2 biological replicates. Dotted line represents the average WT activity. B. Integration efficiency of output library variants, as fold change to WT Dn29. Dots and error bars represent the mean ± SD of n=2 biological replicates. The dotted line represents average WT activity, and gray bands represent the SD of n=36 biological replicates of WT Dn29. C. Integration efficiency of mutations in variant 127 reverted to WT, shown as fold change to variant 127. Bars and error bars represent the mean ± SD of n=2 biological replicates, shown as dots. D. Integration efficiency (orange, left y-axis) and specificity (teal, right y-axis) of variant 127 with position 341 saturation mutagenesis, shown as fold change to WT. Bars and error bars represent the mean ± SD of n=2 biological replicates, shown as dots. E. Integration efficiency (orange, left y-axis) and specificity (teal, right y-axis) of variant 127 with lysine scan mutations of putative DNA binding residues. Bars and error bars represent the mean ± SD of n=2 biological replicates. F. Integration efficiency (orange, left y-axis) and specificity (orange, right y-axis) of variant 381 with significant (one-tailed p<0.05) mutations from second validation round. Bars and error bars represent the mean ± SD of n=6 (variant 381) or n=2 (other variants) biological replicates. G. Integration efficiency vs. specificity of 341K and 341Q lineages with driver mutations. Dots and error bars represent the mean ± SD of n=2 biological replicates for each experimental condition (n=3 for WT and variant 127). H. Epistatic interactions between rounds 2 and 3 mutations. X-axis: expected effect (sum of single mutant log_2_(fold change)); Y-axis: observed effect (double mutant log_2_(fold change)). Dots represent n=2 biological replicates, the dotted line is the identity line. Pearson r=0.2841; two-tailed p=0.0172.

**Extended Data Figure 5:**
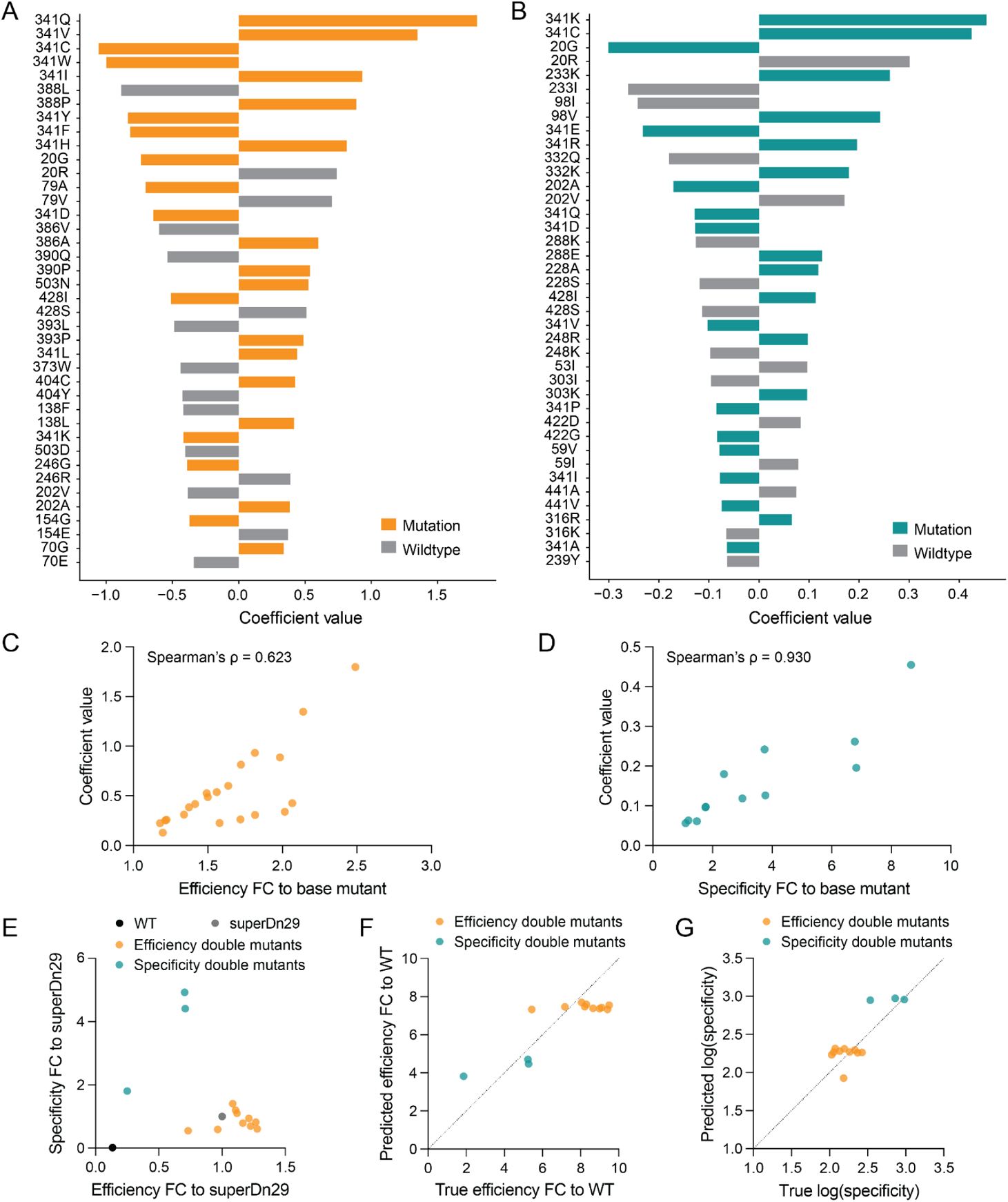
Model coefficients, predictions, and validation for recombinase engineering. A-B. Top 40 coefficients of the (A) efficiency model and (B) specificity model. C-D. Correlation between the (C) efficiency (Spearman’s ρ =0.623, p=0.0025) and (D) specificity (Spearman’s ρ =0.930, p<0.0001) models’ coefficient values and experimental impact of mutations, shown as fold change to the base mutant (variant 127 from mutations identified in the first round of individual validation (Figure 1G, S4C-E), or variant 381 from mutations identified in the second round of individual mutation validation (Figure S4F)). E. Efficiency and specificity of model-guided variants, which each contain two mutations on top of superDn29, and were designed to maximize efficiency (orange) or specificity (teal). Each dot represents the mean of n=2 biological replicates. F-G. Comparison between the model-predicted and true (F) efficiency and (G) specificity of the model-guided variants. Each dot represents the mean of n=2 biological replicates.

**Extended Data Figure 6:**
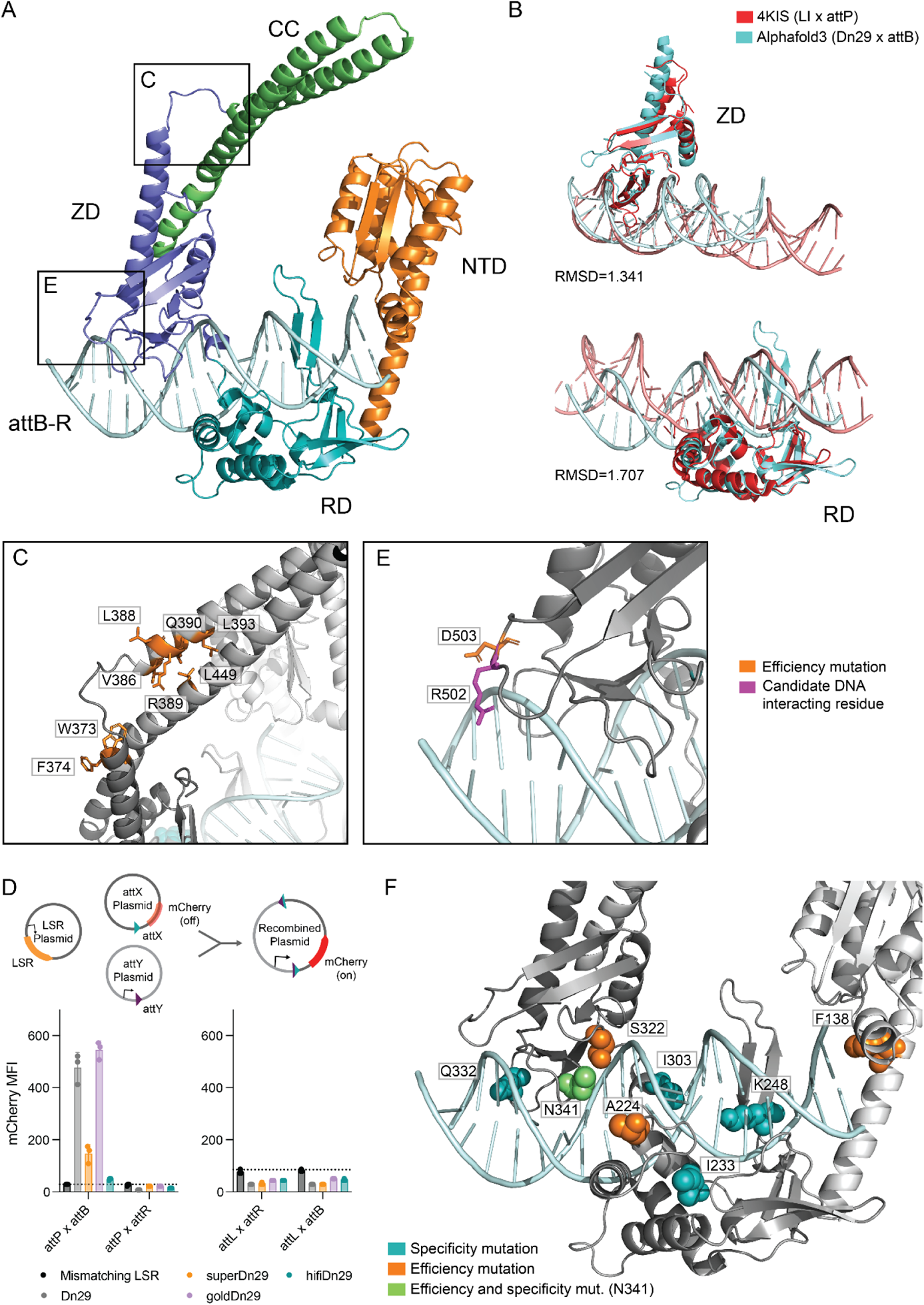
Mapping driver mutations on an Alphafold3 model of Dn29 bound to attB. A. Alphafold3 model of Dn29 bound to the attB-R half site, colored by protein domain. NTD: N-terminal domain, RD: recombinase domain, ZD: zinc-ribbon domain, CC: coiled-coil motif. B. Alignment of the Dn29 x attB-R Alphafold3 structure to listeria integrase (LI) C-terminal domain bound to attP crystal structure (PDB: 4KIS). Top: zinc-ribbon domain (ZD), bottom: recombinase domain (RD). Root mean square deviation (RMSD) values provided. C. Coiled-coil hinge region with efficiency mutations (orange). Corresponds to box C in panel A. D. (Top) schematic of plasmid recombination assay for attachment site recombination. mCherry expresses upon recombination between attachment sites X and Y. (Bottom) recombination of Dn29, key variants, and mismatching LSR control between attP, attB, attL, and attR, measured by mCherry median fluorescence intensity (MFI). Dotted line indicates the background fluorescence associated with the mismatching LSR control. Bars and error bars represent the mean ± SD of n=3 biological replicates, shown as dots. E. Efficiency mutation D503N (orange) and neighboring DNA-interfacing residue R502 (magenta). Corresponds to box D in panel A. F. Specificity (teal) and efficiency (orange) mutations near DNA-interfacing residues. N341 (green) is both a specificity and efficiency mutation.

**Extended Data Figure 7:**
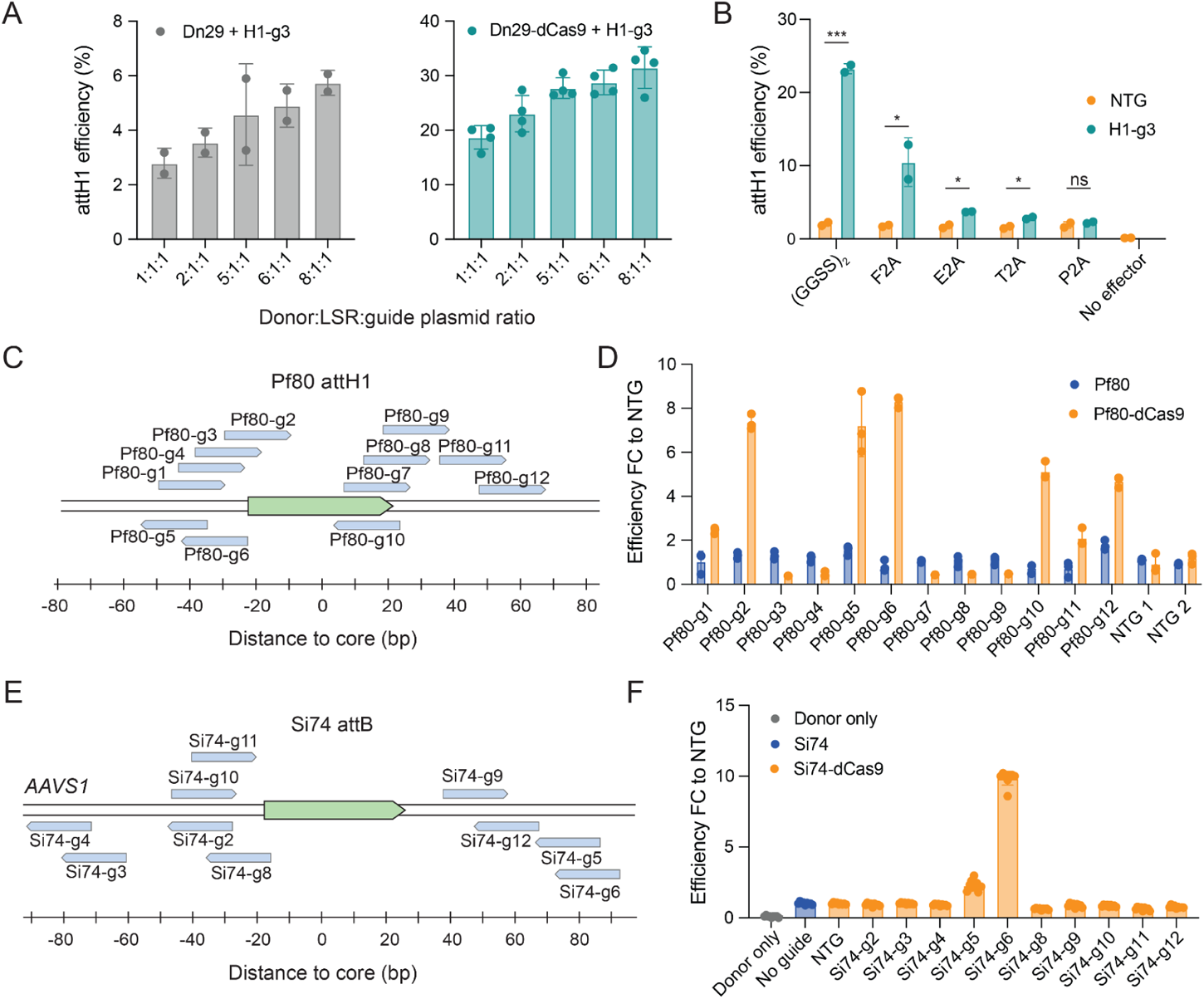
Versatility and optimization of LSR-dCas9 fusions across different recombinases and genomic targets. A. attH1 integration efficiency of donor:effector:guide plasmid stoichiometries for Dn29 (left, n=2) and Dn29-dCas9 (right, n=4). Bars and error bars represent the mean ± SD. B. attH1 integration efficiency of Dn29-dCas9 with direct fusion or 2A peptide linkers, with H1-g3 and non-targeting sgRNA (NTG). 2A peptides ranked in order from least to most complete ribosomal skipping. Bars and error bars represent the mean ± SD of n=2 biological replicates, shown as dots. Asterisks show t-test significance. * = one-tailed p<0.05; ns=not significant. C. Schematic of sgRNA targets for Pf80 attH1 pseudosite (chr11:64,243,293). D. Integration efficiency at Pf80 attH1 pseudosite by Pf80 and Pf80-dCas9, shown as fold change to NTG. Bars and error bars represent the mean ± SD of n=3 biological replicates. E. Schematic of sgRNA targets for Si74 attB, pre-inserted at the *AAVS1* locus. F. Integration efficiency at *AAVS1* Si74 attB with Si74 and Si74-dCas9, shown as fold change to NTG. Bars and error bars represent the mean ± SD of n=3 biological replicates. Dots represent the 3 technical replicates per biological replicate.

**Extended Data Figure 8:**
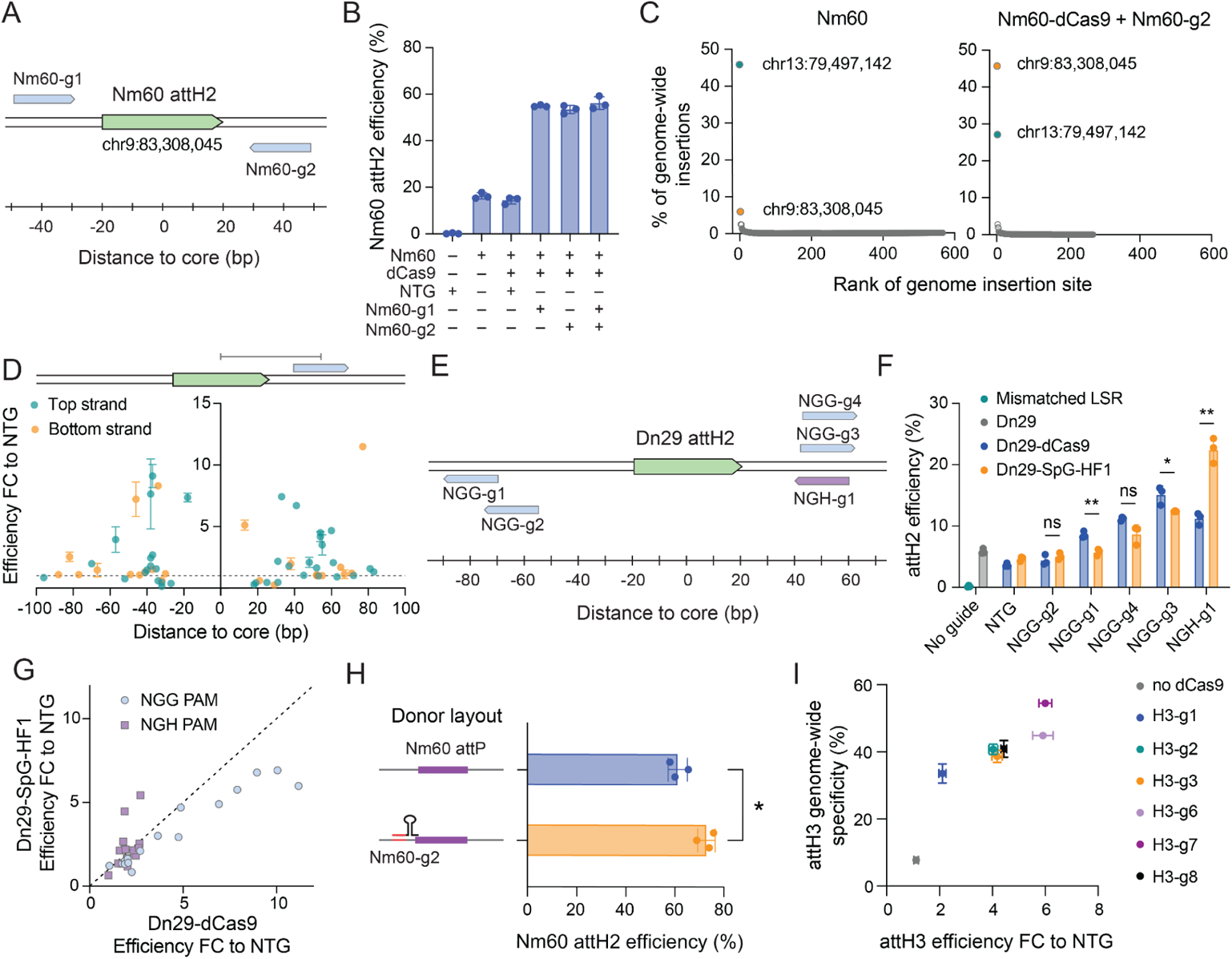
Multiplexing genome sgRNAs and donor sgRNAs with LSR orthologs and dCas9 variants. A. Schematic of sgRNA targets for the Nm60 attH2 pseudosite (chr9:83,308,045). B. Integration efficiency of Nm60-dCas9 at attH2 with guides targeting upstream and downstream of the pseudosite, in single and multiplex. Bars and error bars indicate mean ± SD of n=3 biological replicates. C. Genome-wide specificity of Nm60 and Nm60-dCas9 with Nm60-g2 sgRNA targeting Nm60 attH2. D. Integration efficiency (fold change to NTG) of all LSR-dCas9 fusions (Dn29-dCas9, Pf80-dCas9, Si74-dCas9, Nm60-dCas9) at all pseudosites/gRNAs tested in Figure 3 and Extended Data Figures 7 and 8, relative to the sgRNA distance to the pseudosite core. The distance to the core is measured as the number of bases between the center of the pseudosite core dinucleotide to the center (11th base) of the 23 bp sgRNA target sequence + PAM sequence. The dots and error bars represent the mean ± SD of n=2-6 biological replicates. E. Schematic of sgRNA targets for Dn29 attH2 pseudosite (chr10:58,514,256). Blue targets: NGG PAMs; purple target; NGH PAM. F. Absolute integration efficiency of Dn29-dCas9 compared to Dn29-SpG-HF1 at attH2. Bars and error bars represent the mean ± SD of n=3 biological replicates, shown as dots. Asterisks show t-test significance. * = two-tailed p<0.05; ** = two-tailed p<0.01; ns=not significant. G. Correlation of Dn29-dCas9 vs. Dn29-SpG-HF1 integration efficiency with NGG or NGH PAM sgRNAs. Dots represent the mean of n=3 biological replicates. Dotted line represents the identity line. H. Integration efficiency of Nm60-dCas9 at attH2 with a donor-binding sgRNA (Nm60-g2) plasmid. Bars and error bars represent the mean ± SD of n=3 biological replicates. Asterisks show t-test significance. *=one-tailed p<0.05. I. Correlation between integration efficiency (fold change to NTG) and genome-wide specificity at attH3 of various attH3-targeting sgRNAs. Dots and error bars represent the mean ± SD of n=3 (efficiency) and n=2 (specificity) biological replicates. Data shown is the same as presented in Figure 3E and 3F.

**Extended Data Figure 9:**
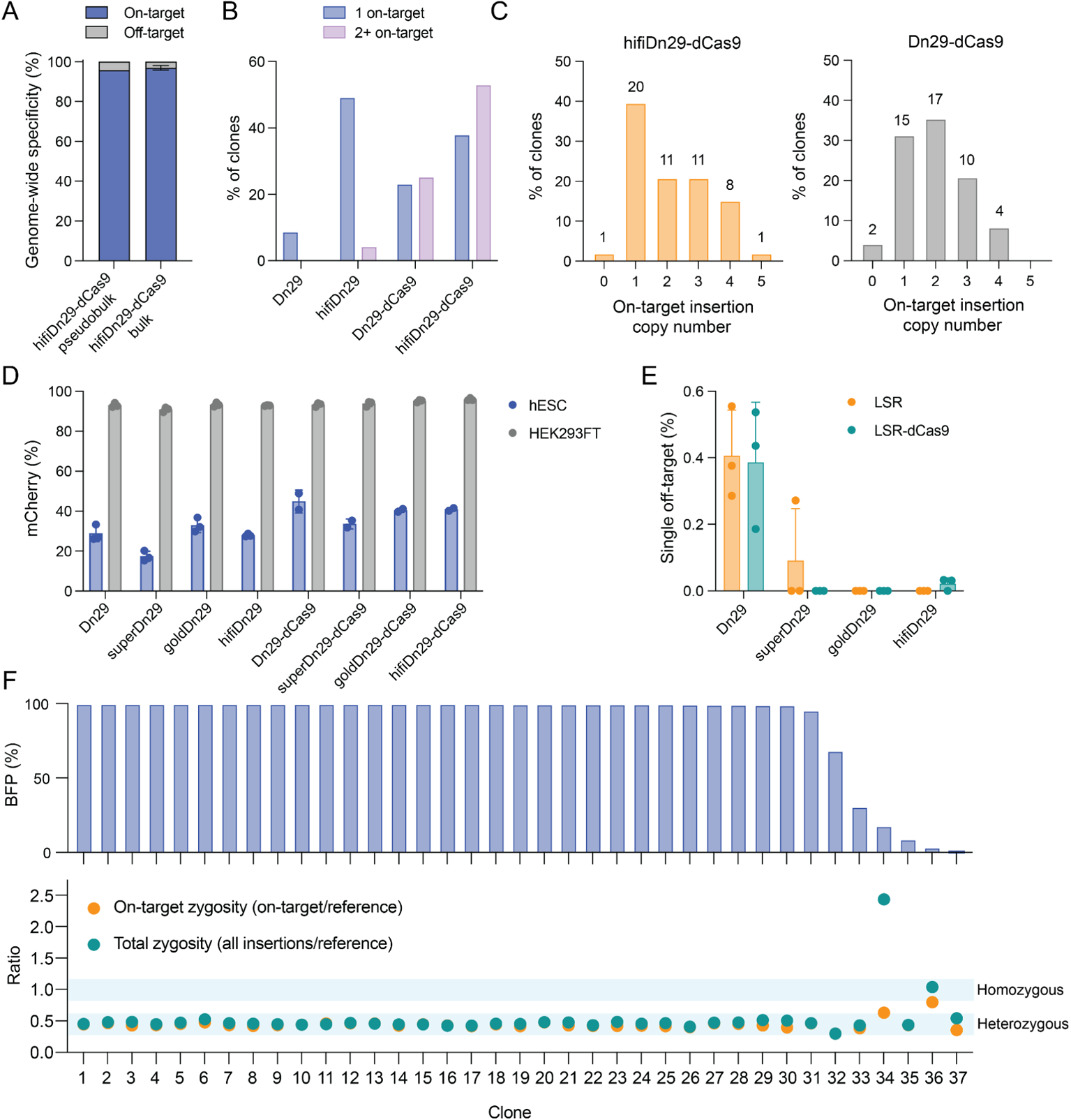
Bulk and single cell characterization of HEK293FTs and H1 hESCs. A. Comparison of hifiDn29-dCas9 on-target and off-target integrations: single cell clonal genotyping (n=53 clones, left) vs bulk genome-wide integration assay (mean ± SD, n=3 biological replicates, right). B. On-target insertion copy number per clone for hifiDn29 and Dn29, with and without dCas9 fusion. C. On-target insertion copy number per clone for hifiDn29-dCas9 and Dn29-dCas9. N of clones is labeled above each bar. D. Donor plasmid transfection efficiency in HEK293FTs and hESCs (% mCherry^+^ cells). Bars and error bars represent the mean ± SD of n=3 biological replicates, shown as dots. E. Specificity of Dn29 and variants in hESCs, measured as attH3 off-target integration efficiency by ddPCR. Bars and error bars represent the mean + SD of n=3 biological replicates, shown as dots. F. H1 hESC clones (n=37) edited with goldDn29-dCas9: BFP expression (top) and genotyping (bottom). Integration/reference ratio of 0.5 indicates heterozygous insertions, 1 indicates homozygous insertions. Single clone per bar/dot.

**Extended Data Figure 10:**
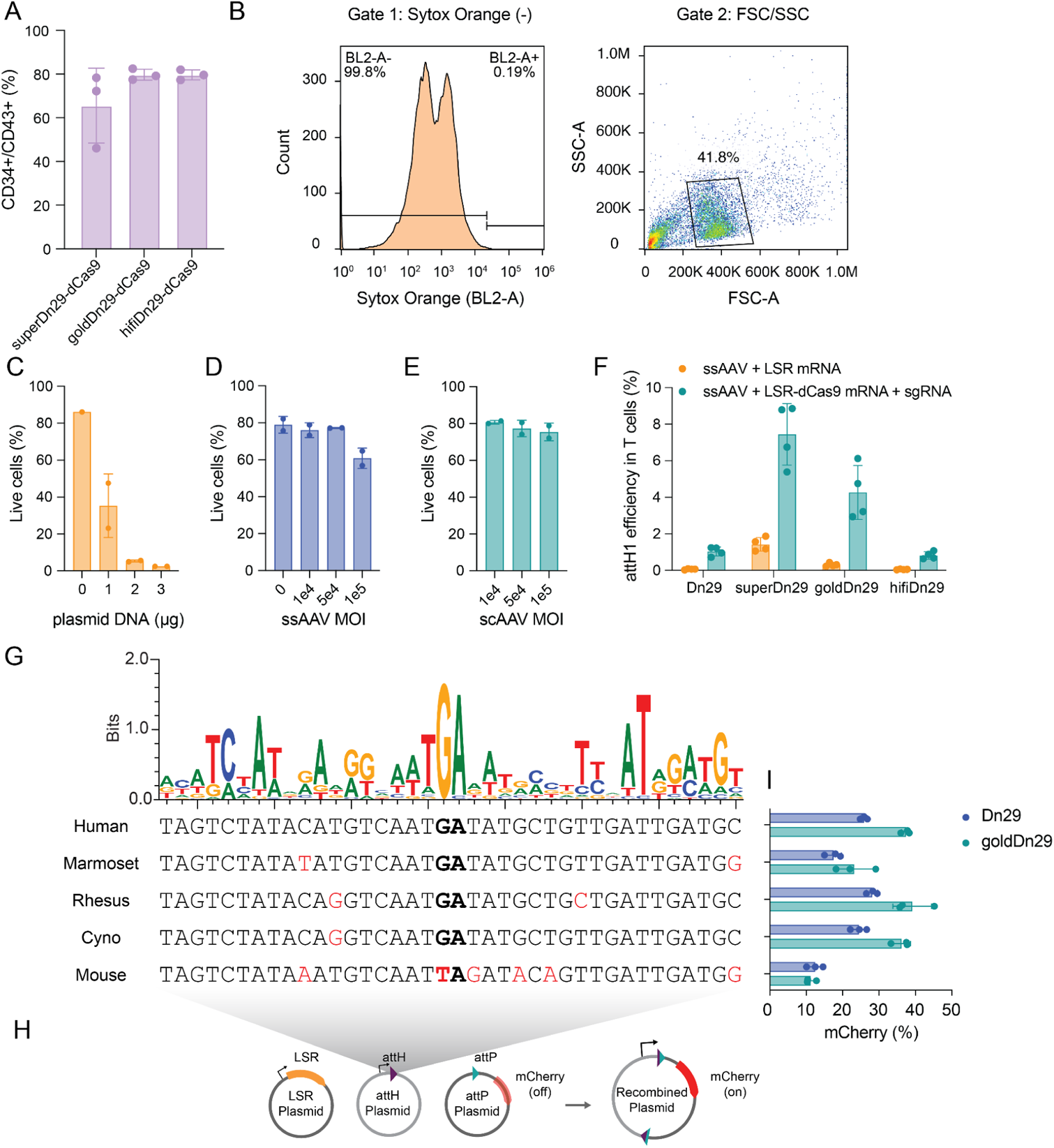
HPC differentiation, T cell engineering, and cross-species compatibility of engineered recombinases. A. HPC differentiation markers (CD34/CD43) of LSR-dCas9-edited hESCs post differentiation. Bars and error bars represent the mean ± SD of n=3 biological replicates. B. Example gating strategy for HPCs. First, unstained cells are used as a negative control to set the Sytox Orange gate, indicating the boundary between live (BL2-A negative) and dead (BL2-A positive) cells (left). Each sample is first gated for Sytox Orange (-), then gated for HPCs using FSC/SSC. Within this population, the median fluorescence intensity (MFI) of the stained cell surface marker is used for determination of knockdown efficiency. C-E. Viability of primary T cells upon (C) electroporation with plasmid DNA, (D) transduction with ssAAV, and (E) transduction with scAAV. Bars and error bars represent the mean + SD of n=2 biological replicates from separate blood donors. F. Integration efficiencies of Dn29 variants and dCas9 fusions at attH1 in primary human T cells using ssAAV donor. Bars and error bars represent the mean ± SD of n=4 biological replicates, each originating from a different blood donor. G. Alignment of attH1-like pseudosites in human, marmoset, rhesus monkey, cynomolgus monkey, mouse and a sequence logo of the top 100 pseudosites in HEK293FTs. H. Schematic of plasmid recombination assay for testing attH1-like pseudosites in HEK293FTs. I. Plasmid recombination efficiency between attP and each pseudosite, using Dn29 and goldDn29, in HEK293FTs. For the mouse pseudosite, the cognate attP plasmid is modified to contain the matching TA dinucleotide core sequence.

## Acknowledgements

We thank Aravind Natarajan, Nora Enright, Ben Mijts, Michael C. Bassik, Hiroshi Nishimasu, Chika Ito, Michael Fanton, Nicholas T. Perry, and all members of the Hsu Lab for helpful discussions; Brian Plosky, Chiara Ricci-Tam, and the Arc Institute Scientific Publications Team for assistance with the manuscript; and the FACS Core and Genomics Platform at Arc Institute for experimental assistance. M.G.D. and S.K. are supported by funding from the Arc Institute. A.F. was partially supported by the NSF Graduate Research Fellowship Program (2019284848). P.D.H. is supported by funding from the Arc Institute, Yosemite, the Biswas Foundation, the Rainwater Foundation, the Curci Foundation, the Rose Hill Innovators Program, S. Altman, V. and N. Khosla, and by anonymous gifts to the Hsu Laboratory.

## Data Availability

The NGS dataset is available on the NCBI Sequence Read Archive at Bioproject PRJNA1172311.

## Author Contributions

A.F. and P.D.H. conceived the study. A.F., L.J.B., and P.D.H. designed experiments. A.F., L.J.B., J.Q.M., V.Q.T., L.G. and J.W. performed experiments. A.F., M.G.D., L.J.B., and V.Q.T. performed computational analyses. A.F., L.J.B., V.Q.T., J.Q.M., L.G., J.W. and P.D.H. analyzed and interpreted the data. P.D.H. provided overall supervision of the research, with assistance from S.K., A.M., and L.A.G.. A.F., A.P., and P.D.H. wrote the manuscript with input from all authors.

